# Century-scale methylome stability in a recently diverged *Arabidopsis thaliana* lineage

**DOI:** 10.1101/009225

**Authors:** Jörg Hagmann, Claude Becker, Jonas Müller, Oliver Stegle, Rhonda C. Meyer, Korbinian Schneeberger, Joffrey Fitz, Thomas Altmann, Joy Bergelson, Karsten Borgwardt, Detlef Weigel

## Abstract

There has been much excitement about the possibility that exposure to specific environments can induce an ecological memory in the form of whole-sale, genome-wide epigenetic changes that are maintained over many generations. In the model plant *Arabidopsis thaliana*, numerous heritable DNA methylation differences have been identified in greenhouse-grown isogenic lines, but it remains unknown how natural, highly variable environments affect the rate and spectrum of such changes. Here we present detailed methylome analyses in a geographically dispersed *A. thaliana* population that constitutes a collection of near-isogenic lines, diverged for at least a century from a common ancestor. We observed little DNA methylation divergence whole-genome wide. Nonetheless, methylome variation largely reflected genetic distance, and was in many aspects similar to that of lines raised in uniform conditions. Thus, even when plants are grown in varying and diverse natural sites, genome-wide epigenetic variation accumulates in a clock-like manner, and epigenetic divergence thus parallels the pattern of genome-wide DNA sequence divergence.

## INTRODUCTION

Differences in DNA methylation between individuals can be due to genetic variation, stochastic events or environmental factors. Epigenetic marks such as DNA methylation are not randomly distributed across plant genomes, but associate with certain classes of genomic loci, especially with transposable elements (TEs). Changes in the DNA sequence or structure caused by, for instance, TE insertion, can induce secondary epigenetic effects at the concerned locus [1,2], or, via RdDM, even at distant loci [3–5]. The high degree of sequence variation, including insertions/deletions (indels), copy number variants (CNVs) and rearrangements among natural accessions in *A. thaliana* provides ample opportunities for linked epigenetic variation [6–10]. The genomes of *A. thaliana* accessions from around the globe are rife with differentially methylated regions (DMRs) [10], but it remains unclear how many of these cannot be explained by closely linked genetic mutations and how many are pure epimutations [11] that occur in the absence of any genetic differences.

The seemingly spontaneous occurrence of heritable DNA methylation differences has been documented for wild-type *Arabidopsis thaliana* isogenic lines grown for several years in a stable greenhouse environment [12,13]. Truly spontaneous switches in methylation state are most likely the consequence of incorrect replication or erroneous establishment of the methylation pattern during DNA replication [14–16]. A potential amplifier of stochastic noise is the complex and diverse population of small RNAs that are at the core of RNA-directed DNA methylation (RdDM) [17] and that serve as epigenetic memory between generations. The exact composition of small RNAs at silenced loci can vary considerably between individuals [13], and stochastic inter-individual variation has been invoked to explain differences in remethylation, either after development-dependent or induced demethylation of the genome [18,19]. Such epigenetic variants can contribute to phenotypic variation within species, and epigenetic variation in otherwise isogenic individuals has been shown to affect ecologically relevant phenotypes in *A. thaliana* [20–22].

In addition to these spontaneous epigenetic changes the environment can induce demethylation or de novo methylation in plants, for example after pathogen attack [23]. Recently, it has been proposed that repeated exposure to specific environmental conditions can lead to epigenetic differences that can also be transmitted across generations, constituting a form of ecological memory [24–27]. The responsiveness of the epigenome to external stimuli and its putative memory effect have moved it also into the focus of attention for epidemiological and chronic disease studies in animals [28,29]. How the rate of trans-generational reversion among induced epivariants with phenotypic effects compares to the strength of natural selection, which in turn determines whether natural selection can affect the population frequency of epivariants, is largely unknown [30–33].

To assess whether a variable and fluctuating environment is likely to have long-lasting effects in the absence of large-scale genetic variation, we have analyzed a lineage of recently diverged *A. thaliana* accessions collected across North America. Using a new technique for the identification of differential methylation, we found that in a population of thirteen accessions originating from eight different locations and diverged for more than one hundred generations, only 3% of the methylome had undergone a change in methylation state. Epimutations at the DNA methylation level did not accumulate at higher rates in the wild as they did in a benign greenhouse environment. Using genetic mutations as a timer, we demonstrate that accumulation of methylation differences was non-linear, corroborating our previous hypothesis that shifts in methylation states are generally only partially stable and that reversions to the initial state are frequent [12,34]. Many methylation variants that segregated in the natural North American lineage could also be detected in the greenhouse-grown population, indicating that similar forces determined spontaneous methylation variation, independently of environment and genetic background. Population structure could be inferred from differences in methylation states, and the pairwise degree of methylation polymorphism was linked to the degree of genetic distance. Together, these results suggest that the environment makes only a small contribution to trans-generationally inherited epigenetic variation on a genome-wide scale.

## RESULTS

### Characterization of the near-isogenic HPG1 lineage from North America

Previous studies of isogenic mutation accumulation (MA) lines raised in uniform greenhouse conditions identified many apparently spontaneously occurring pure epimutations [12,13]. To determine whether variable and fluctuating environments in the absence of large-scale genetic variation substantially alter the genome-wide DNA methylation landscape in the long term, we analyzed a lineage of recently diverged *A. thaliana* accessions collected across North America. Different from the native range of the species in Eurasia, about half of all North American individuals appear to be identical when genotyped at 139 genome-wide markers [35]. We selected 13 individuals of this lineage, called haplogroup-1 (HPG1), from locations in Michigan, Illinois and on Long Island, including pairs from four sites (Figure 1a, Table S1). Whole-genome sequencing of pools of eight to ten siblings from each accession identified a shared set of 670,979 single nucleotide polymorphisms (SNPs) and 170,998 structural variants (SVs) relative to the Col-0 reference genome, which were then used to build a HPG1 pseudo reference genome (SOM: Genome analysis of HPG1 individuals; Table S2; Figure S1).

**Figure 1:**
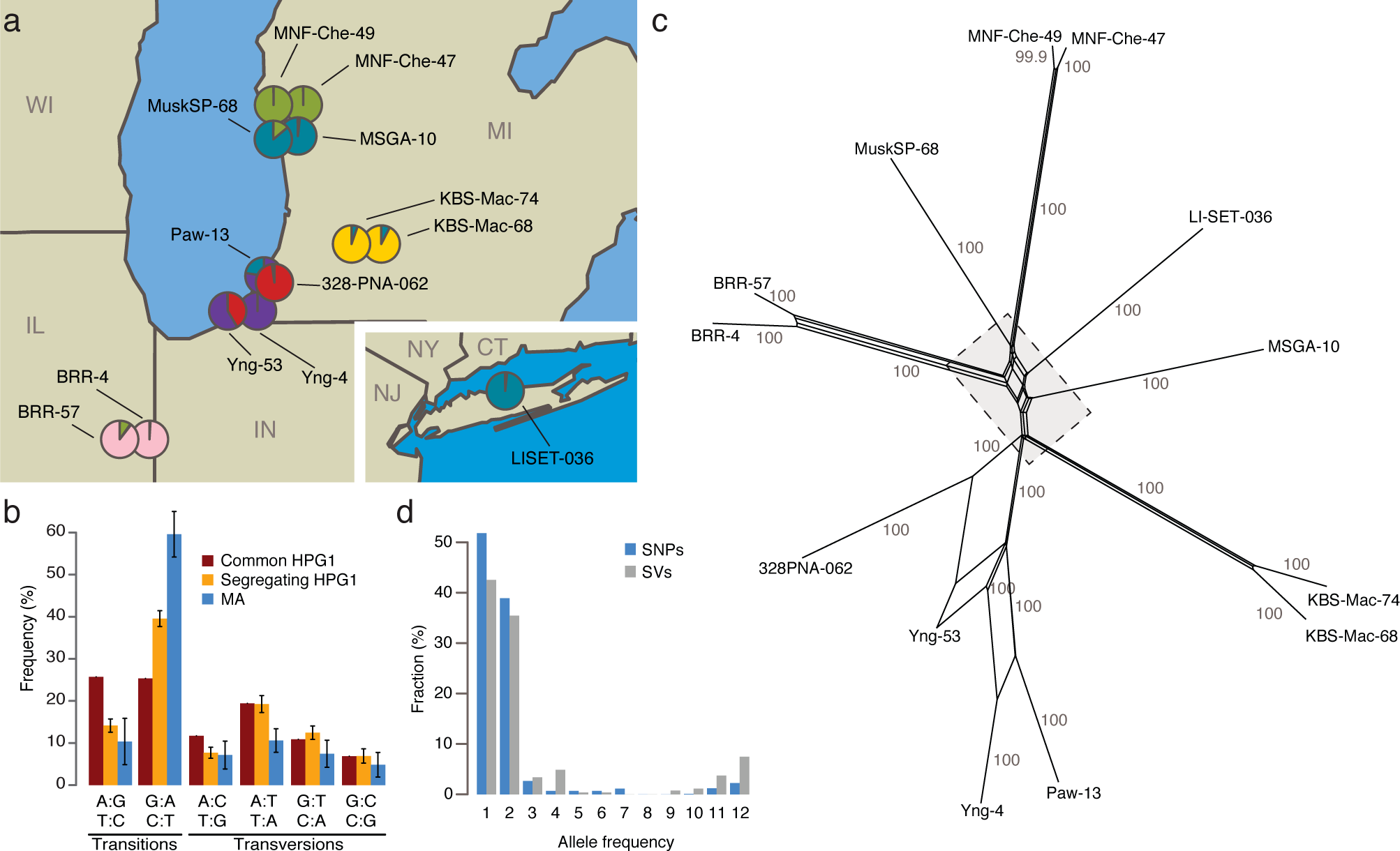
North American HPG1 accessions belong to a genetically homogeneous population. (a) Sampling locations of the 13 strains analyzed in this study. Pie charts indicate population structure inferred from segregating SNP data. Data were analyzed using STRUCTURE[60], with *K*=6. CT=Connecticut, IL=Illinois, IN=Indiana, MI=Michigan, NJ=New Jersey, NY=New York, WI=Wisconsin. (b) Single-nucleotide mutation spectrum. Bars represent the accession average, error bars indicate 95% confidence intervals. (c) Phylogenetic network of HPG1 accessions based on segregating SNPs and SVs with SplitsTree v.4.12.3 [62]. Numbers indicate bootstrap confidence values (10,000 iterations). Dashed line delimits close-up in Figure S4. (d) Allele frequencies of SNPs.

Only 1,354 SNPs and 521 SVs segregated in this population (Table S3, Figure S2 and S3), confirming that the 13 strains were indeed closely related. Segregating SNPs were noticeably more strongly biased towards GC→AT transitions than shared SNPs, especially in TEs, although the bias was not as extreme as in the greenhouse-grown MA lines (Figure 1b) [36]. A phylogenetic network and STRUCTURE analysis based on the segregating polymorphisms reflected the geographic origin of the accessions (Figure 1a, c; Figure S4). Three of the pairs of accessions from the same site were closely related, and were responsible for many alleles with a frequency of 2 in the sampled population (Figure 1d). If the spontaneous genetic mutation rate is similar to that seen in the greenhouse [36], the HPG1 accessions would be 15 to 384 generations separated from each other. With a generation time of one year, their most recent common ancestor would have lived about two centuries ago, which is consistent with *A. thaliana* having been introduced to North America during colonization by European settlers [37]. Lastly, we observed only a weak positive correlation between genetic distance and phenotypic difference (Figure S5). We conclude that the HPG1 accessions constitute a near-isogenic population that should be ideal for the study of heritable epigenetic variants that arise in the absence of large-scale genetic change under natural growth conditions.

**Figure 2:**
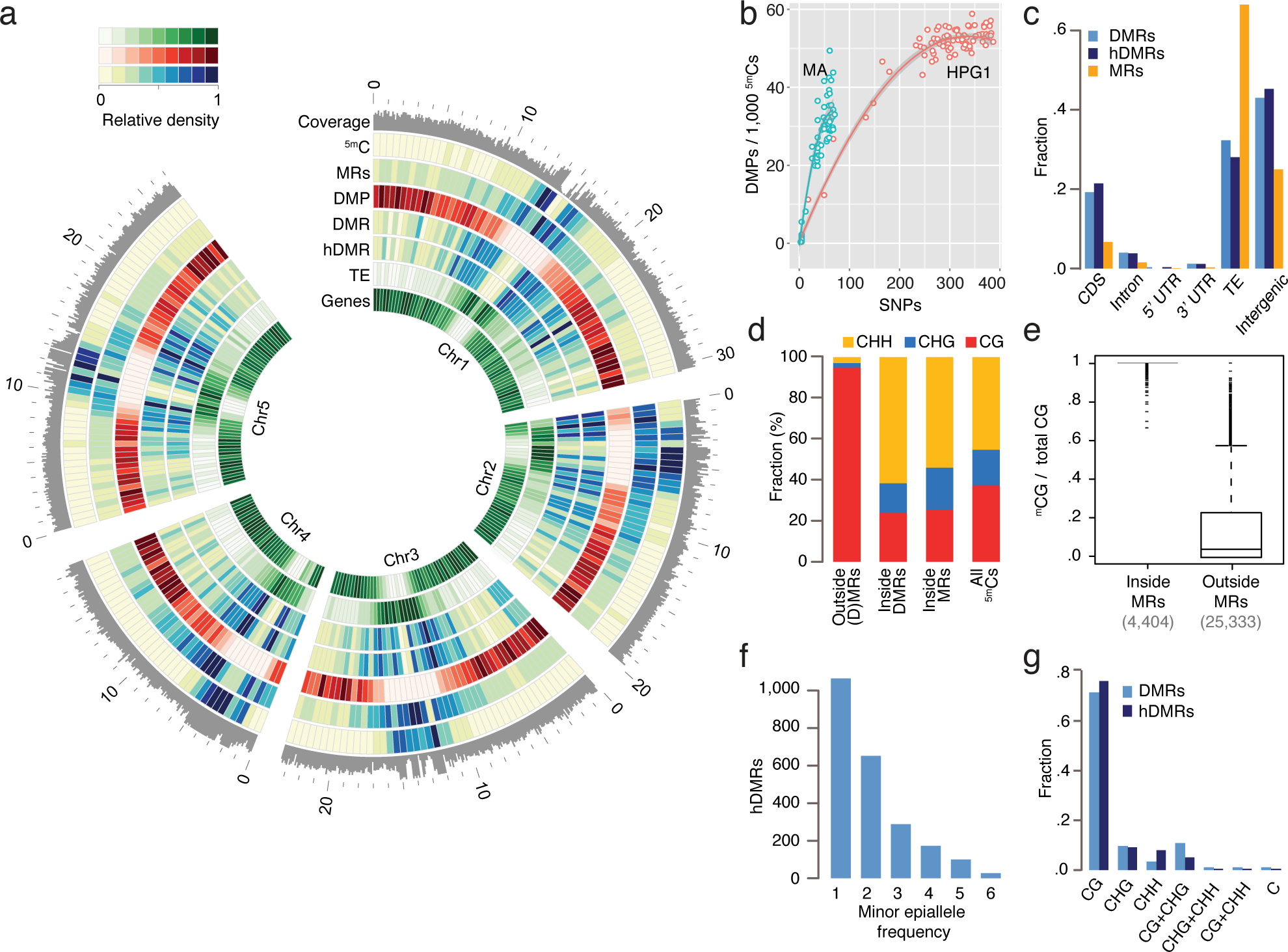
Epigenetic variation in a nearly isogenic population. (a) Genome-wide features: average coverage in 100 kb windows, the remainder in 500 kb windows. Outside coordinates in Mb. (b) DMP number in relation to SNP number in pairwise comparisons. MA data are based on single individuals, HPG1 data on pools of 8-10 individuals; each data point represents an independent comparison of two lines. DMPs in each pairwise contrast were scaled to the number of methylated sites compared. (c) Annotation of cytosines in MRs and hDMRs. (hD)MR sequences were assigned to only one annotation in the following order: CDS > intron > UTR > transposon > intergenic. (d) Sequence context of methylated positions relative to MRs and DMRs. (e) Fraction of ^5m^CGs among all CG sites for each gene and transposable element, with at least 5 CGs. (f) Minor epiallele frequencies of 2,304 hDMRs that could be split into only two groups and for which at least four strains showed statistically significant differential methylation. Strains not tested statistically significant for a particular hDMR were not considered for this plot. (g) DMRs and hDMRs according to sequence contexts in which significant methylation differences were found. ‘C’ denotes (h)DMRs in all three contexts.

**Figure 3:**
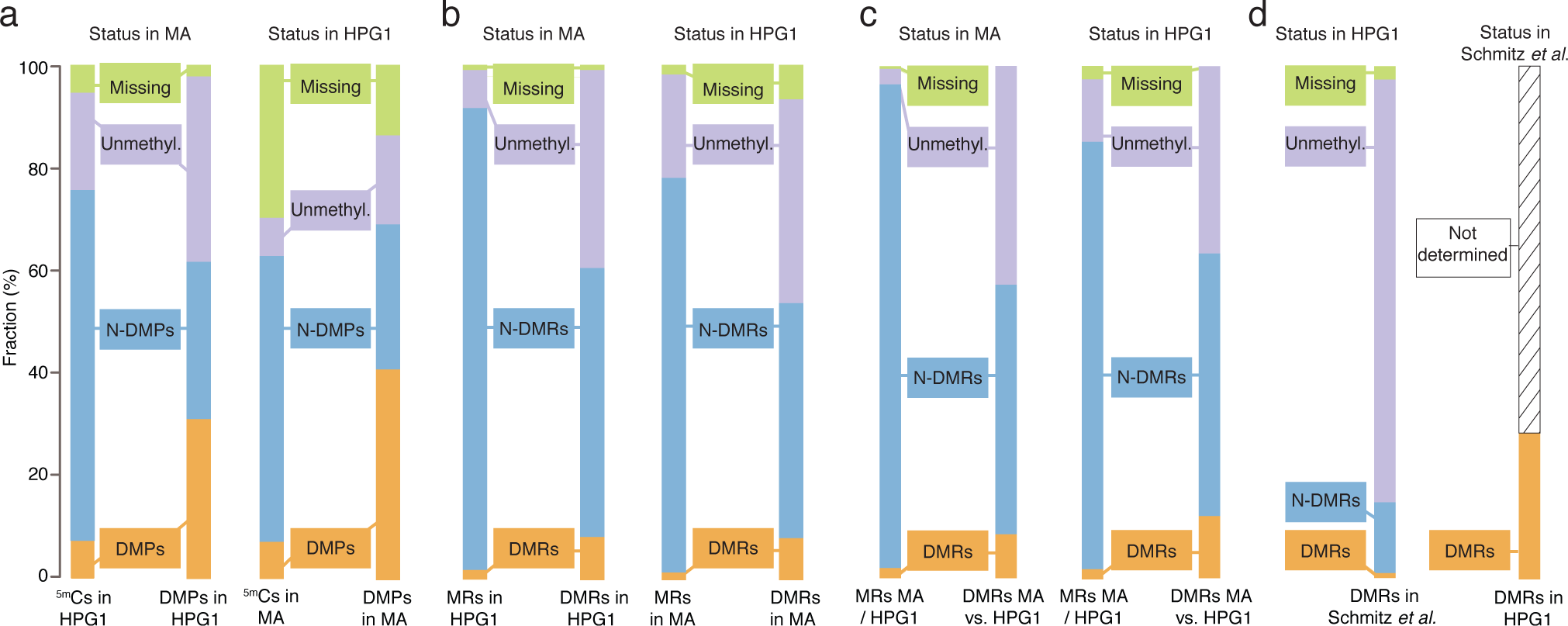
Epigenetic variation in independent populations and conditions. (a) Comparison of ^5m^Cs and DMPs identified in pairwise comparisons of HPG1 and MA lines^12^. Left: sites in HPG1 strains and their status in the MA data; right: sites in the MA strains and their status in the HPG1 data. (b) Comparison of MRs and DMRs identified in pairwise comparisons of HPG1 and MA lines. Left: regions in HPG1 strains and their status in the MA data; right: regions in the MA strains and their status in the HPG1 data. (c) MRs and DMRs identified in comparison between one randomly chosen MA line (30-39) and one randomly chosen HPG1 line (MuskSP-68), and their overlap with within-population DMRs. (d) Comparison of HPG1 DMRs with DMRs identified in 140 natural *A. thaliana* accessions [10]. Because MRs were not reported in ref. [10], the overlap of DMRs with non-DMRs could not be assessed.

**Figure 4:**
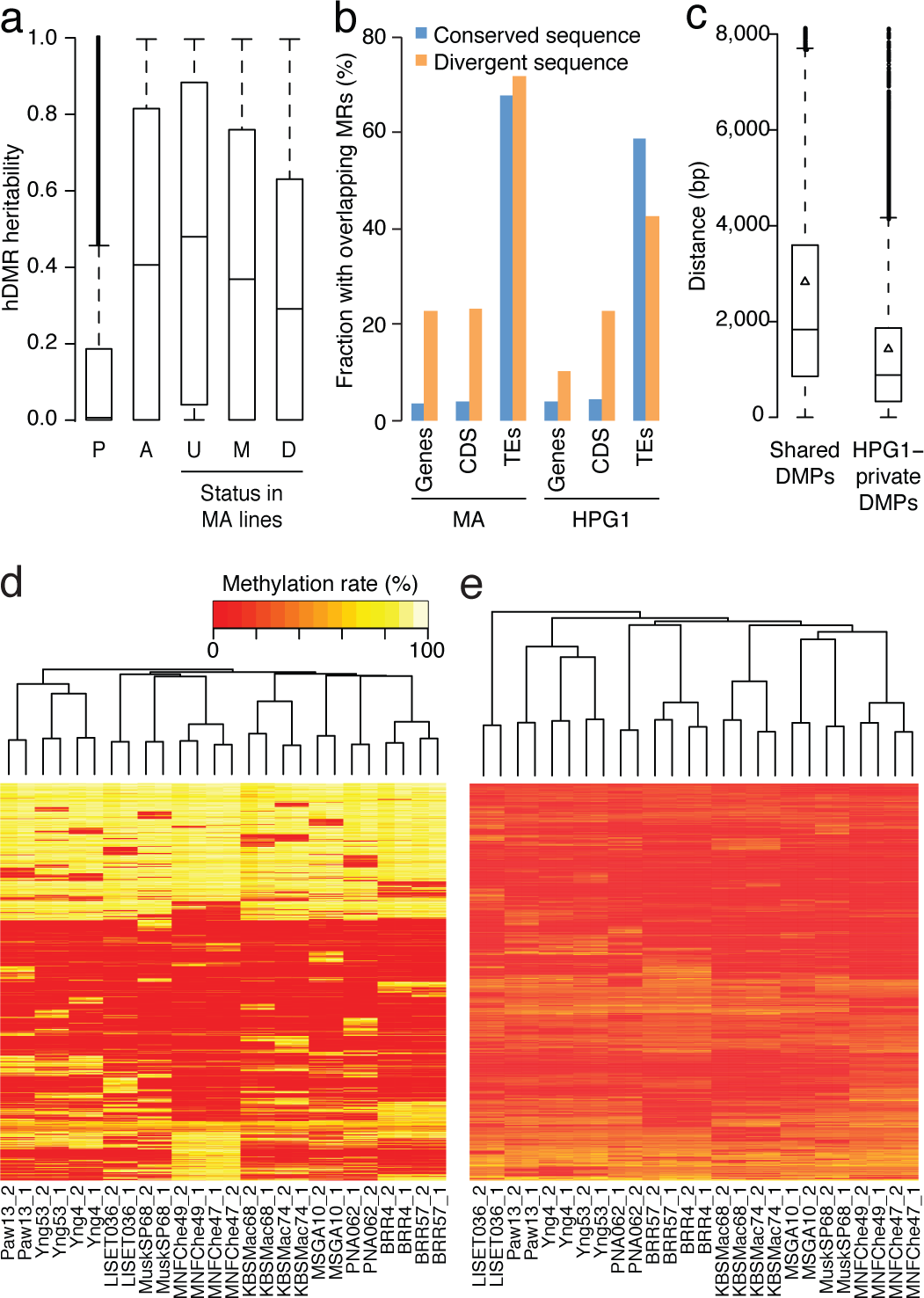
Genetic effects on epigenetic variation. (a) Heritability values based on genome-wide genetic differentiation for all hDMRs, hDMRs with randomly permuted methylation rates and subsets of hDMRs depending on their overlap with MA-MRs and MA-DMRs. P: Permuted (2,945 hDMRs); A: All (2,945); U: Unmethylated in MA (1,310); M: Methylated in MA (1,243); D: DMR in MA (392). (b) Correlation between SVs and probability of overlap with MRs. Divergent sequences are insertions of at least 20 bp relative to the other population. This analysis is based on 3,256 SVs overlapping with genes, 641 with CDS and 4,020 with TEs (Table S11). (c) Distances between common SVs of at least 20 bp and the closest DMP, depending on whether it is shared between the MA and HPG1 populations. Triangles represent the mean. (d) Hierarchical clustering of HPG1 strains based on methylation rates at 50,000 CG-DMPs. (e) Hierarchical clustering of HPG1 strains based on average methylation rates of 2,829 hDMRs with full information across all strains. Methylation rates per region were calculated as the average methylation rate of each methylated cytosine in that region.

### Differentially methylated positions the HPG1 lineage

To assess the long-term heritable fraction of DNA methylation polymorphisms in the HPG1 lineage, we grew plants under controlled conditions for two generations after collection at the natural sites, before performing whole methylome bisulfite sequencing on two pools of 8-10 individuals per accession (SOM: Primary analysis of methylation; Table S4). After mapping reads to the HPG1 pseudo reference genome, we first investigated epigenetic variation at the single-cytosine level. There were 535,483 unique differentially methylated positions (DMPs), with an average of 147,975 DMPs between any pair of accessions (SD = 23,745); thus, 86% of methylated cytosines accessible to our analyses were stably methylated across all HPG1 accessions. The vast majority of DMPs (97%) was detected in the CG context (CG-DMPs). As we have discussed previously [12], this can be largely attributed to the differences in methylation rates. Because of lower average CHG and CHH methylation compared to CG methylation at individual sites, statistical tests of differential methylation fail more often for CHG and CHH sites. The finding that only about 2% of all covered cytosines were differentially methylated strongly contrasts with a previous population epigenomic study [10], which despite lower sequencing depth than our experiments concluded that the vast majority, over 90%, of all cytosines in the *A. thaliana* genome is differentially methylated among 140 natural, more divergent accessions [10], with a third being found at minor allele frequencies of over 10%.

Using the geographic outlier LISET-036 as a reference strain, we found that 61% of CG-DMPs as well as 36% of the small number of CHG- and CHH-DMPs were present in at least two independent accessions (Figure S6a), many of them shared between accessions from the same site. As is typical for *A. thaliana* [38], most methylated positions clustered around the centromere and localized to TEs and intergenic regions (Figure 2a; Figure S6b). In contrast, CG-DMPs were over-represented on chromosome arms, localizing predominantly to coding sequences (Figure 2a; Figure S6b), similar to what we had previously observed in the greenhouse-grown MA lines [12].

We asked whether DMPs had accumulated more quickly in natural environments than in the greenhouse, using DNA mutations in the HPG1 and MA populations as a molecular clock (SOM: Estimating DMP accumulation rates). Our null hypothesis was that a variable and highly fluctuating natural environment increases the rate of heritable methylation changes. In contrast, DMPs accumulated in sub-linear fashion in both the HPG1 and MA populations [12] (Figure 2b) – with similar trends for DMPs in all three contexts – and DMPs did not accumulate more rapidly in the HPG1 than in the MA lines. The steeper initial increase relative to SNP differences as well as the broader distribution of MA line differences relative to HPG1 differences were most likely the result of having compared individual plants in the MA experiment [12], rather than pools of siblings, as in the HPG1 experiment (Figure S7; SOM: Estimating DMP accumulation rates). Furthermore, if the genetic mutation rate in the wild were higher than in the greenhouse, for example because of increased UV exposure, we would underestimate the epimutation rate per generation in the HPG1 strains.

### Differentially methylated regions the HPG1 lineage

Because it is unclear what consequences variation at individual methylated cytosines has in plants, we next investigated differentially methylated regions (DMRs) in the HPG1 population. A limitation of previous plant methylome studies using short read sequencing has been that these relied on DMPs or fixed sliding windows along each chromosome to identify DMRs, rather than beginning with what appears intuitively to be more appropriate, namely regions that are known to be methylated in individual strains (methylated regions, MRs; SOM: Methylated regions) [39]. We therefore adapted a Hidden Markov Model (HMM), which had been developed for segmentation of animal methylation data [40], to the more complex DNA methylation patterns in plants (SOM: Differentially methylated regions). We identified on average 32,529 MRs per strain (median length 122 bp), with almost a quarter of the HPG1 reference genome, 22.6 Mb, covered by an MR in at least one strain (Figure 2a, c; Table S5, Figure S8a). MRs overlapping with coding regions were over-represented in genes responsible for basic cellular processes (p-value << 0.001), in agreement with gene body methylation being a hallmark of constitutively expressed genes [41]. Only 1% of ^m^CHH and 2% of ^m^CHG positions were outside MRs (Figure 2d), consistent with the dense CHH and CHG methylation found in repeats and silenced TEs [38]. Compared to ^m^CGs within MRs, ^m^CGs outside MRs localized almost exclusively to genes (94%), were spaced much farther apart, and were separated by many more unmethylated loci (Figure 2e; Figure S8b, c). This explains why sparsely methylated genes were under-represented in HMM-determined MRs, even though gene body methylation accounts for a large fraction of ^m^CGs. The accuracy of our MR detection method was well supported by independent methods (SOM: Differentially methylated regions).

Using the unified set of MRs, we tested all pairs of accessions for differential methylation, identifying 4,821 DMRs with an average length of 159 bp. Of the total genome space occupied by MRs, only 3% were contained in DMRs, indicating that the heritable methylation patterns had remained largely stable in this set of geographically dispersed accessions (Figure S8a, e; Figure S9; Table S6). Indeed, 91% of genic and 98% of the TE sequence space were devoid of DMRs. Of the DMRs, 3,199 were classified as highly differentially methylated (hDMRs; Table S7). Their allele frequency spectrum was similar to that of DMPs (Figure 2f). Most DMRs and hDMRs showed highly variable methylation in only one cytosine context, often CG (Figure 2g). Different from DMPs, the densities for DMRs and hDMRs were highest in centromeric and pericentromeric regions, and overlapped more often with TEs than with genes (Figure 2a, c). In relation to the full complement of MRs, however, genic regions were two-fold overrepresented in the genome sequence covered by DMRs, and three-fold in the genome sequence covered by hDMRs (Figure 2c). In a recent report of 140 divergent accessions [10], DMRs were also biased towards genic regions, but not quite as extreme as in the HPG1 lines, likely reflecting the much greater genetic variation among TEs in this set of accessions [10], compared to the only recently diverged HPG1 lines.

### Methylation variation and transcriptome changes

DNA methylation in gene bodies has been proposed to exclude H2A.Z deposition and thereby stabilize gene expression levels [41]. We therefore asked what impact differential methylation had on transcriptional activity. We identified 269 differentially expressed genes across all possible pairwise comparisons (Table S8, S9), most of which were found in more than one comparison. When we clustered accessions by differentially expressed genes, closely related pairs were placed together (Figure S11). We identified 28 differentially expressed genes that overlapped with an hDMR either in their coding or 1 kb upstream region, but the relationship between methylation and expression was variable (Table S10). By visual examination of hDMRs, we found not more than five instances of demethylation that were associated with increased expression; examples are shown in Figure S12.

### Comparison between genetic and epigenetic differentiation

With the caveat that there are uncertainties about the genetic mutation rate in the wild, and therefore how the number of SNPs relates to the number of generations since the last common ancestor, there was no evidence for faster accumulation of DMPs in the HPG1 population, nor for very different epimutation rates among HPG1 lines (Figure 2b). Importantly, the overlap between DMPs in the two populations was much greater than expected by chance: the chance of a random ^m^C site in the MA population of being a DMP in the HPG1 population was only 7%, but it was 41% among sites that were also DMPs in the MA population. In other words, compared to all ^m^C sites in the MA population, DMPs in the HPG1 population were four-to six-fold enriched among sites that were also DMPs in the MA population, and vice versa (Figure 3a) (SOM: Similarity of epigenetic variation profiles in independent populations). Shared DMPs were more heavily biased towards the chromosome arms and towards genic sequences than population-specific DMPs (Figure S13a and S13b). Conversely, DMPs from one population were more likely to be unmethylated throughout the other population when compared to random methylated sites (Figure 3a), as one might expect for sites that sporadically gain methylation.

DMPs private to the HPG1 lineage appeared to be less frequent in the pericentromere compared to DMPs private to the MA lines (Figure S13a), which was also reflected in an apparently higher epimutation frequency in the MA lines for these regions (Figure S13b). We therefore investigated whether the annotation spectrum differed between these two classes of DMPs. Even though MA-private DMPs were more often found in TEs compared to HPG1-private DMPs, this bias was also observed for all cytosines accessible to our methylome analyses (Figure S13c), and can therefore be explained by a more accurate read mapping and better TE annotation in the Col-0 reference compared to the HPG1 pseudo-reference genome. Indeed, except for chromosome 4, the average sequencing depth in the pericentromere was higher in the MA lines (Figure S13b).

DMPs distinguishing MA lines that were separated by only a few generations more frequently overlapped with HPG1 DMPs than DMPs identified between distant MA lines (Figure S14). We interpret this observation as an indication of privileged sites that are more labile and therefore more likely to have changed in status already after a small number of generations.

Similar to variable single positions, or DMPs, the overlap between 2,523 DMRs in the MA lines and the 4,821 DMRs of the HPG1 accessions was highly significant (Z-score = 32.9; 100,000 permutations) (Figure 3b). We observed similar degrees of overlap independently of DMR sequence context. Overlapping DMRs were, in contrast to shared DMPs, not biased towards genic regions (Figure S15). DMRs of the HPG1 lineage, however, overlapped with genic sequences more often than MA-DMRs (Figure S15), which might again be explained by the different efficiencies in mapping to repetitive sequences and TEs (Figure S13b). We identified DMRs that distinguish the MA and HPG1 populations using a randomly chosen MA and a randomly chosen HPG1 line; these DMRs, which differentiate distantly related accessions, were also enriched in each of the two sets of within-population DMRs (MA or HPG1) (Figure 3c). Finally, we compared HPG1-DMRs to DMRs that had been identified with a different method among 140 natural accessions from the global range of the species[10] (Figure 3d). Although only 9,994, less than one fifth, of the DMRs from the global accessions were covered by methylated regions in the HPG1 strains, the overlap of DMRs was highly significant (Z-score = 19.8; 100,000 permutations). In conclusion, the high recurrence of DMPs and DMRs from different datasets points to the same loci being inherently biased towards undergoing changes in DNA methylation independently of genetic background and growth environment.

To quantify how many methylation differences were co-segregating with genome-wide genetic changes in cis and trans, we estimated heritability for each hDMR by applying a linear mixed model-based method. We used segregating sequence variants with complete information as genotypic data and average methylation rates of hDMRs with complete information as phenotypes. The median heritability of all hDMRs was 0.41 (mean 0.44), which means that genetic variance across the entire genome contributed less than half of methylation variance (Figure 4a). Regions classified as hDMRs in the HPG1 strains that were not methylated in the greenhouse-grown MA lines had a higher median heritability, 0.48, than HPG1 hDMRs also found among MA DMRs (0.29), which held true for all sequence contexts (Figure 4a; Figure S16). hDMRs found only in the HPG1 population, especially those in unmethylated regions of the MA lines, were thus more likely to be linked to whole-genome genotype than hDMRs found in both populations. For 19% of all hDMRs (21% CG-hDMRs, 14% CHG-hDMRs, 7% CHH-hDMRs), the whole-genome genotype explained more than 90% of their methylation differences (with a standard error of at most 0.1). Of these hDMRs, half had a heritability of greater than 0.99. That 6.7% of the sequence space of these heritable hDMRs still overlapped with MA DMRs (versus 9.4% for the less heritable hDMRs) was in agreement with the hypothesis that there are regions that vary highly in their methylation status independently of genetic background.

To identify genetic variants that potentially directly cause methylation changes in their local genomic neighborhood, we focused on DMRs with segregating SNPs or indels located within 1 kb. Of 191 such DMRs, only three showed a systematic correlation with nearby sequence polymorphisms. We noticed, however, that coding regions with SVs larger than 20 bp that distinguished the MA and HPG1 populations were more likely to be methylated in both the MA and HPG1 lines than non-polymorphic coding regions (Figure 4b). Consequently, HPG1-specific DMPs were on average closer to SVs than DMPs shared between the HPG1 and MA populations (Figure 4c).

Next, we asked whether the genome-wide methylation pattern reflected genetic relatedness, i.e., population structure. Hierarchical clustering by methylation rates of DMPs and hDMRs grouped strains by sampling location (Figure 4d, e). This result was largely independent of the sequence or the annotation context of the DMPs and hDMRs, and not seen with N-DMPs (Figure S17). That MRs not classified as DMRs (N-DMRs) grouped the accessions similar to DMPs, albeit with less confidence (shorter branch lengths; Figure S17), suggested that our DMR calling algorithm was conservative. Methylation data thus paralleled similarity between accessions at the genetic level, in agreement with methylation differences reflecting the number of generations since the last common ancestor.

## DISCUSSION

We have tested the hypothesis that under natural conditions, epigenetic variation accumulates over the short term in a manner that is very different from the clock-like behavior of genetic variation [24–27], by taking advantage of a unique natural experiment, a lineage that has likely diverged for at least a century throughout North America. Our analyses have revealed little evidence for long-term heritable genome-wide epigenetic differentiation that might have been induced by the variable and fluctuating environmental conditions experienced by the HPG1 accessions since they separated from each other. While the exact conditions these plants have been subjected to since their separation remain unknown, the time scale and diversity of geographic provenance are strong indicators of the variability of the environment between the different sampling sites. The general framework enabled by the HPG1 lineage – nearly isogenic lines grown for more than a century under variable and fluctuating conditions – could not have been achieved in a controlled greenhouse experiment.

Studies of epiRIL populations have shown that pure epialleles can be stably transmitted across generations [5,19], but how often this is the case for environmentally induced epigenetic changes has been heavily debated [33,42–44]. The recent excitement about the transmission of induced epigenetic variants comes from such induced variants having been proposed to be more often adaptive than random genetic mutations [25–27]. Contrary to the expectations discussed above, we found that epimutation rates under natural growth conditions at different sites did not exceed those observed in a controlled greenhouse environment, with polymorphisms accumulating sub-linearly in both situations, apparently because of frequent reversions. Note that we grew the HPG1 plants under controlled conditions for two generations after sampling at the natural site, to reduce the range of epigenetic variation to the long-term heritable fraction. We cannot exclude that in field-grown HPG1 individuals epigenetic variation is increased and carries a stronger signature of the sampling site. However, such a hypothetical fraction of epigenetic variation, if it existed, is not heritable, because we did not find evidence for it after two extra generations in the greenhouse. Additional studies comparing plants grown outdoors to their progeny grown in a stable and controlled environment will help to further clarify these issues.

That DMPs between closely related MA lines are more likely to overlap with HPG1 DMPs than DMPs between more distantly related MA lines supports the hypothesis that there are different classes of polymorphic sites. One of these includes ‘high lability’ sites that are independent of the genetic background, that change with a high epimutation rate, and that are therefore more likely to appear in each population. Another class of DMPs comprises more stable sites that gain or lose methylation more slowly and that therefore are less likely to be shared between different populations.

Differences between accessions in terms of DNA methylation recapitulated their genetic relatedness, further corroborating our hypothesis that heritable epigenetic variants arise predominantly as a function of time rather than as a consequence of rapid local adaptation. Epigenetic divergence thus does not become uncoupled from genetic divergence when plants grow in varying environments, nor does the rate of epimutation increase. A minor fraction of heritable epigenetic variants may be related to habitat, which could be responsible for LISET-036 being epigenetically a slight outlier (Figure 4e), even though it is not any more genetically diverged from the most recent common ancestor of HPG1 than other lines. Such local epigenetic footprints could also explain fluctuations in epimutation frequency between the MA and HPG1 lineages. Subtle adaptive changes at a limited number of loci would go unnoticed in the present analysis of genome-wide patterns and can therefore not be excluded. However, on a genome-wide scale there was little indication of adaptive change: neither were LISET-036 specific DMRs in and near genes enriched for GO terms with an obvious connection to environmental adaptation, nor were there overlapping differentially expressed genes (Figure S18, SOM: Analysis of LISET-036 specific hDMRs). In combination with the general lack of correlation between differential methylation and changes in gene expression, our findings suggest that epigenetic changes in nature are mostly neutral, and thus comparable to genetic mutations.

Because of the near-isogenic background of the HPG1 accessions, we were also able to gauge how much of epigenetic variation is either caused by, or stably co-segregates with genetic differences. HPG1-specific hDMRs were more often linked to genotype variation than regions that were variably methylated in both the HPG1 and MA populations. This suggests that heritable hDMRs can, to a certain extent, be considered facilitated epigenetic changes [11].

Even though DMRs, like DMPs, are over-represented in genes, they are mainly located in TEs and intergenic regions, which is different from the situation for DMPs. Altogether our data indicate that both DMPs and constitutively methylated sites in genes are typically separated by many unmethylated sites and that a large fraction is therefore not classified as being within (D)MRs. Variability of DNA methylation in genes thus mainly affects single, sparsely distributed cytosines. Further experiments are necessary to clarify the biological relevance of variation of single-site DNA methylation in genic regions.

In summary, comparisons between MA laboratory strains and natural HPG1 accessions revealed that DMPs overlapped much more than expected by chance, despite these populations having experienced very different environments that also differ greatly in their stability, and despite completely different genetic backgrounds. The observation that changes at many sites and loci are independent of the genetic background and geographic provenance suggests that spontaneous switches in methylation predominantly reflect intrinsic properties of the DNA methylation and gene silencing machinery. Our most important finding is probably that DNA methylation is highly stable across dozens, if not hundreds of generations of growth in natural habitats; 97% of the total methylated genome space was not contained in a DMR. This is in stark contrast to published data describing more than 90% of the genome as variably methylated in a set of 140 divergent natural accessions [10]. We propose that heritable polymorphisms that arise in response to specific growth conditions appear to be much less frequent than those that arise spontaneously. These conclusions are of importance when considering epimutations as a potential evolutionary force.

## MATERIAL AND METHODS

### Plant growth and material

Accessions [35] were collected in the field at locations indicated in Table S1. Seeds had been bulked in the Bergelson lab at the University of Chicago before starting the experiment. Plants were then grown at the Max Planck Institute in Tübingen on soil in long-day conditions (23 °C, 16 h light, 8 h dark) after seeds had been stratified at 4 °C for 6 days in short-day conditions (8 h light, 16 h dark). We grew one plant of each accession under these conditions; seeds of that parental plant were then used for all experiments. Eight plants of the same accession were grown per pot in a randomized setup. All accessions used in this paper have been added to the 1001 Genomes project (http://1001genomes.org) and have been submitted to the stock center.

### Nucleic acid extraction

DNA was extracted from rosette leaves pooled from eight to ten individual adult plants. Plant material was flash-frozen in liquid nitrogen and ground in a mortar. The ground tissue was resuspended in Nuclei Extraction Buffer (10 mM Tris-HCl pH 9.5, 10 mM EDTA, 100 mM KCl, 0.5 M sucrose, 0.1 mM spermine, 0.4 mM spermidine, 0.1% b-mercaptoethanol). After cell lysis in nuclei extraction buffer containing 10% Triton-X-100, nuclei were pelleted by centrifugation at 2000 *g* for 120 s. Genomic DNA was extracted using the Qiagen Plant DNeasy kit (Qiagen GmbH, Hilden, Germany). Total RNA was extracted from rosette leaves pooled from eight to ten individual adult plants using the Qiagen Plant RNeasy Kit (Qiagen GmbH, Hilden, Germany). Residual DNA was eliminated by DNaseI (Thermo Fisher Scientific, Waltham, MA, USA) treatment.

### Library preparation

DNA libraries for genomic and bisulfite sequencing were generated as described previously [12]. Libraries for RNA sequencing were prepared from 1 µg of total RNA using the TruSeq RNA sample prep kit from Illumina (Illumina) according to the manufacturer’s protocol.

### Sequencing

All sequencing was performed on an Illumina GAII instrument. Genomic and bisulfite-converted libraries were sequenced with 2 x 101 bp paired-end reads. For bisulfite sequencing, conventional *A. thaliana* DNA genomic libraries were analyzed in control lanes. Transcriptome libraries were sequenced with 101 bp single end reads. Four libraries with different indexing adapters were pooled in one lane; no control lane was used. For image analysis and base calling, we used the Illumina OLB software version 1.8.

### Processing of genomic reads

The SHORE pipeline v0.9.0 [45] was used to trim and quality-filter the reads. Reads with more than 2 (or 5) bases in the first 12 (or 25) positions with a base quality score of less than 4 were discarded. Reads were trimmed to the right-most occurrence of two adjacent bases with quality values equal to or greater than 5. Trimmed reads shorter than 40 bases and reads with more than 10% (of the read length) of ambiguous bases were discarded.

### Genetic variant identification

Genetic variants were called in an iterative approach. In each step, SNPs and structural variants common to all strains were detected and incorporated into a new reference genome. The thus refined “HPG1-like” genomes served as the reference sequence in the subsequent iterations (Figure S1). We performed three iterations to call segregating variants and built two reference genomes to retrieve common polymorphisms. The steps performed in each iteration will be described in the following.

### Read mapping

Reads were aligned against the *Arabidopsis thaliana* genome sequence version TAIR9 in iteration 1 and against updated “Haplogroup 1-like” genomes in further iterations. The mapping tool GenomeMapper v0.4.5s [46] was used, allowing for up to 10% mismatches and 7% single-base-pair gaps along the read length to achieve high coverage. All alignments with the least amount of mismatches for each read were reported. A paired-end correction method was applied to discard repetitive reads by comparing the distance between reads and their partner to the average distance between all read pairs. Reads with abnormal distances (differing by more than two standard deviations) were removed if there was at least one other alignment of this read in a concordant distance to its partner. The command line arguments used for SHORE are listed in Supplementary File 1.

### SNP and small indel calling

Base counts on all positions were retrieved by SHORE v0.9.0 [45] and a score was assigned to each site and variant (SNP or small indel of up to 7% of read length) depending on different sequence and alignment-related features. Each feature was compared to three different empirical thresholds associated with three different penalties (40%, 20% and 5% reduction of the score, initial score: 40). They can be found in Table S12.

For comparisons across lines, positions were accepted if at most one intermediate penalty on their score was applicable to at least one strain (score **≥** 32). In this case, the threshold for the other strains was lowered, accepting at most one high and two intermediate penalties (score **≥** 15). In this way, information from other strains was used to assess sites from the focal strain under the assumption of mostly conserved variation, allowing the analysis of additional sites.

Only sites sufficiently covered (**≥** 5x) and with accepted base calls in at least half of all strains (**≥** 7 out of 13) were processed further. Variable alleles with a frequency of 100% were classified as "common" and variants with a lower frequency as "segregating".

Additional SNPs were called using the targeted *de novo* assembly approach described below.

### Structural variant (SV) calling

Although a plethora of SV detection tools have been developed, their predicted SV sets show little overlap between each other on the same data sets. Furthermore, the false positive rate of many methods can be drastic [47]. Hence, rather than taking the intersection of the output from different tools, which would yield only a low amount of SVs, we combined different tools and applied a stringent evaluation routine to identify as many true SVs as possible. Since SVs common to all strains should be incorporated into a new reference, only methods that identify SVs on a base pair level could be used. Currently, there are four different SV detection strategies (based on depth of coverage, paired-end mapping, split read alignments or short read assembly, respectively). Only tools based on split read alignments and assemblies are capable of pinpointing SV breakpoints down to the exact base pair. Programs that were used include Pindel v2.4t [48], DELLY v0.0.9 [49], SV-M v0.1 [50] and a custom local *de novo* assembly pipeline targeted towards sequencing gaps (described below). We reported deletions and insertions from all tools, and additionally inversions from Pindel. DELLY combines split read alignments with the identification of discordant paired-end mappings. Thus, our SV calling made use of three out of four currently available methodologies.

Reads for DELLY were mapped using BWA v0.6.2 [51] against the TAIR9 Col-0 reference genome to produce a BAM file as DELLY’s input format.

The arguments for the command line calls of all tools are listed in Supplementary File 1.

### Targeted *de novo* assembly

While using a re-sequencing strategy, there are regions without read coverage (“sequencing gaps”) because either the underlying sequence is being deleted in the newly sequenced strain, or highly divergent to the reference sequence, or present in the focal strain, but not represented in the read set. To access the strain’s sequences of the first two cases, a local *de novo* assembly method was developed.

Insertion breakpoints or small deletions, however, can mostly not be detected by zero coverage due to reads ranging with a few base pairs into or beyond the structural variants. Therefore, we defined a “core read region” as the read sequence without the first and last 10 nucleotides. To be able to assemble the latter cases, the definition of “sequencing gaps” was extended from zero-covered regions to stretches not spanned by a single read’s core region.

All reads aligned to the surrounding 100 nucleotides of such newly defined sequencing gaps as well as the unmappable reads from the re-sequencing approach together with their potential mapped partners constituted the assembly read set. Two assembly tools were used to generate contigs, SOAPdenovo2 v2.04 [52] and Velvet v1.2.0 [53] (command line arguments in Supplementary File 1). Contigs shorter than 200 bp were discarded. To map the remaining contigs of each assembler against the iteration-specific reference genome, their first and last 100 bp were aligned with GenomeMapper v0.4.5s [46], accepting a maximal edit distance of 10. If both contig ends mapped uniquely within 5,000 bp, the thus framed region on the reference was aligned to the contig using a global sequence alignment algorithm after Needleman-Wunsch (‘needle’ from the EMBOSS v6.3.1 package). In addition, non-mapping contigs were aligned with blastn (from the BLAST v2.2.23 package) [54] to yield even more variants.

All differences between contig and reference sequences were parsed (including SNPs, small indels and SVs) for each assembly tool. Only identical variants retrieved from both assemblers were selected.

### Generating and filtering consolidated variant set of each strain

For each strain, all variants from the SV tools and the *de novo* assemblies were consolidated (Figure S1a) and positioned to consistent locations to be comparable using the tool Dindel v1.01 [55]. In the case of contradicting or overlapping variants, identical variants (having the same coordinates and length after re-positioning) predicted by a majority of tools were chosen and the rest discarded, or all were discarded if there was no majority.

Despite sequencing errors or cross-mapping artifacts of the re-sequencing approach, genomic regions covered by reads are generally trusted. Chances of long-range variations in the inner 50% of a mapped read’s sequence (“inner core region” of a read) are assumed to be low, since gaps would deteriorate the alignment capability towards the ends of the read.

Therefore, we filtered out variants from the consolidated variant set spanning a genomic region already covered by at least one inner core region of a mapped read of the corresponding strain (Figure S1a), assuming homozygosity throughout the genome. This “core read criterion” had to be fulfilled at each genomic position spanned by the variant.

### Using branched reference to validate variants

Variants passing the core read filter in all strains were classified as common variants and were incorporated into the reference sequence of the previous iteration, thus replacing the reference allele. Segregating variants, which could not be detected in all strains, were additionally built into the reference in separate “haplotype regions” (or “branches” of the reference sequence) to eventually be able to assess whether reads preferentially mapped to the reference or the alternative haplotype sequence (Figure S1a). Linked variant haplotypes of a strain (distance between consecutive variants **≤**107 bp, the maximal possible span of a read on the reference) as well as identical haplotype regions among strains were merged into one branch sequence.

For each strain, all reads were re-mapped to this new reference sequence yielding read counts at the variant site on each branch (r_b_) and at the corresponding site on the reference haplotype sequence (r_ref_) (Figure S1a). Here, the read count of a site was defined as the number of inner core regions spanning the site. To increase certainty of variant calling and to rule out heterozygosity, the read count of the major allele was tested against a binomial distribution that assumed 95% allele frequency out of a total of r_b_+r_ref_ observations, i.e. sole presence of either the branch or the reference haplotype (if 100% had been assumed, it would not yield a distribution). The null hypothesis of homozygosity was rejected after *P* value correction by Storey’s method [56] for *q* values below 0.05.

The same variant could be part of several different haplotypes and thus, could be included into different branch sequences. Reads supporting this variant would map at multiple locations in the reference. Therefore, we allowed all aligned rather than only unique reads to contribute to read counts and omitted the paired-end correction procedure.

### Final sets of common and segregating variants

We followed a similar “population-aware” approach to prefer commonalities among strains as was used for the SNP calling for labeling variants as being common or segregating. Here, variable sites with accumulated coverage over both branch and reference sequence of **≤** 3× were marked as “missing data”. If there was at least one haplotype in a strain with a *q* value above 0.05, it was assumed to be present in the population. If the test on the same haplotype failed in another strain, but the absolute read count of the haplotype sequence exceeded the alternative haplotype read count by **≥** 2-fold, then this haplotype was considered present in the corresponding strain as well.

We classified variants where at least 7 out of 13 strains did not show missing data as ‘common’ if the branched haplotype was present in all strains, as ‘not present’ if the reference haplotype was present in all strains, or into ‘segregating’ if there was support for both haplotypes.

To combine common variants identified by the described stepwise algorithm into potentially less evolutionary events, we aligned 200 bp around each variant of the last iteration’s genome back to the TAIR9 Col-0 reference genome using a global alignment strategy (‘needle’ from the EMBOSS v6.3.1 package).

In total, we found 842,103 common and 2,017 segregating polymorphisms without removing linked loci compared to Col-0 after two iterations, to which the different tools contributed to different extent depending on the variant type (Figure S1c).

### Methylome sequencing

Genomic and bisulfite sequencing were performed as described in ref. [12].

### Processing and alignment of bisulfite-treated reads

The procedure followed one described [12], except that we aligned reads against the HPG1-like as well as against the Col-0 reference genome sequences. Command line arguments for SHORE are listed in Supplementary File 1.

### Determination of methylated sites

We followed the same procedures as described [12]. Here, we restricted the set of analyzed positions to cytosine sites with a minimum coverage of 3 reads and sufficient quality score (Q25) in at least half of all strains (i.e. **≥** 7), that is, 21 million positions in total. Out of those, we identified 3.8 million methylated cytosines in at least one strain by applying a false discovery rate (FDR) threshold at 5%, and between 2,120,310 and 2,927,447 methylated sites per strain (Table S4). False methylation rates retrieved from read mapping against the chloroplast sequence can be found in Table S4.

### Identification of differentially methylated positions (DMPs)

We performed the same methods as in Becker *et al*. to obtain DMPs[12]. First, cytosine positions were tested for statistical difference between both replicates of a sample using Fisher’s exact test and a 5% FDR threshold. Because individual samples consisted of a pool of several plants, the number of DMPs between replicates was negligible (between 0 and 161). After excluding them, we applied Fisher’s exact test on the 3.8 million cytosine sites methylated in at least one strain for all pairwise strain comparisons. Using the same *P* value correction scheme as in Becker *et al*., we identified 535,483 DMPs across all 13 strains.

### Identification of methylated regions (MRs)

To statistically detect stretches of positions consistently methylated higher as their flanking regions, we used a Hidden Markov Model (HMM) implementation modified from Molaro and colleagues [40]. It assumes that the number of methylation-supporting reads at each cytosine follows a beta binomial distribution and that distributions over positions within and between methylated regions will differ from each other, providing a way to distinguish them. To do so, the HMM uses two states for high and low methylation. The method of Molaro and colleagues was designed for calling MRs in human samples, where the vast majority of methylated cytosines are in a CG context. In plants, however, one observes considerable methylation in all three contexts (CG, CHG and CHH), each with a different methylation rate distribution. Hence, we extended the HMM by learning the parameters of three different beta binomial distributions per state, one for each context. Additionally, in contrast to humans, only the minority of cytosines in the CG context is methylated, as are cytosines in the other contexts. Hence, methylation rates were inverted to find hypermethylated, rather than hypomethylated regions as in the original HMM implementation.

Apart from these changes, we followed the same computational steps as described by Molaro and colleagues [40]: The describing parameters of the – in our case – six distributions (determining the emission probabilities) and the transition probabilities between states were iteratively trained (using the Baum-Welch algorithm) from methylation rates of all cytosines in the corresponding context throughout the genome. After each iteration, all cytosines were probabilistically classified into the most likely state via Posterior Decoding, given the trained model. After training of the HMM, i.e. after maximally 30 iterations or when convergence criteria were met, consecutive stretches of high methylation state were scored, in our case by the sum of all contained methylation rates. Next, *P* values were computed by testing the scores against an empirical distribution of scores obtained by random permutation of all cytosines throughout the genome. After FDR calculation, consecutive stretches in high state with an FDR < 0.05 are defined as methylated regions (MRs).

The HMM was run on all genome-wide cytosines, independent of their coverage. Methylation rates were obtained using accumulated read counts from the strain replicates, resulting in one segmentation of the genome per strain. Gaps of at least 50 bp without a covered C position within a high methylation state automatically led to the end of the high methylation segment. Positions with a methylation rate below 10% at the start or end of highly methylated regions (until the first position with a rate larger than 10%), were assigned to the preceding or subsequent low methylation region, respectively.

### Identification of differentially methylated regions (DMRs)

The method to identify MRs yielded 13 different segmentations of the genome, one for each strain. We selected regions being in different or highly methylated states between strains and statistically tested them for differential methylation (including FDR calculation) as described below. To obtain epiallele frequencies, we clustered strains into groups based on their pairwise comparisons and statistically tested the groupings against each other. Regions that showed statistically significant methylation differences between at least two sets of strains were identified as DMRs. Finally, because of the sensitivity of the statistical test, we empirically filtered DMRs for strong signals and defined highly differentially methylated regions (hDMRs).

### Selecting regions to test for differential methylation

We defined a breakpoint set containing the start and end coordinates of all predicted methylated regions. Each combination of coordinates in this set defined a segment to perform the test for differential methylation in all pairwise comparisons of the strains, if at least one strain was in a high methylation state throughout this whole segment (Figure S9a). To also detect quantitative differences rather than solely presence/absence methylation, we also compared entirely methylated regions in more than one strain to each other.

Because of the sheer number of such regions, we applied the following greedy filter criteria: Regions were discarded from any pairwise comparison if less than 2 strains contained at least 10 cytosines covered by at least 3 reads each (accumulated over strain replicates) in this region (Figure S9a (a)). Regions were discarded from any pairwise comparison if the reciprocal overlap of this region to at least one previously tested region was more than or equal to 70% (Figure S9a (b)). This was done to prevent “similar” regions to be tested twice. Pairwise tests of a region were not performed if both strains were in low methylation state throughout the whole region (Figure S9a (c)). Strains were excluded from pairwise comparisons in a region if the number of positions covered by at least 3 reads each was less than half of the maximum number of such positions of all strains in the same region (Figure S9a (d)). This prevented comparing regions with unbalanced coverage to each other, e.g. a strain with 10 data points against another one with only 2.

These filters reduced the set of regions to test from ∼2.5 million to ∼230,000 per pairwise comparison.

### Testing regions for differential methylation between strains

We designed a statistical test for differential methylation between two strains for a given region. The test assumes that the number of methylated and unmethylated read counts per position along a region follows a beta binomial distribution – similar to the HMM in MR calling. More precisely, there are 3 distributions for each sequence context and for each strain. Using gradient-based numerical maximum likelihood optimization, we fitted the parameters for each beta binomial distribution on the available read count data of the region in the respective strain. This was done a) for each of the two strains separately (while taking strain replicates into account), resulting in (two times three) *strain-specific* beta binomial distributions, and b) for the read counts of both strains including their replicates together, resulting in (three) *common* beta binomial distributions. In this way, we obtained each distribution’s mean μ and standard deviation σ. We selected only regions for potential DMRs, whose intervals [μ_1_ – 2σ_1_, µ_1_ + 2σ_1_] for strain 1 and [µ_2_ – 2σ_2_, µ_2_ + 2σ_2_] for strain 2 did not overlap.

To further corroborate statistical significance, we computed *P* values by calculating the ratio of the *strain-specific* and the *common* log likelihoods of the available read count data using the corresponding beta binomial distributions and by testing it against a chi-squared distribution (with 6 degrees of freedom). Let sample *S* have *N*_Sc_ cytosines in context *c* in total and *C_Scp_* reads at position *p* in context *c*, from which *x_Scp_* are methylated, then we compute:

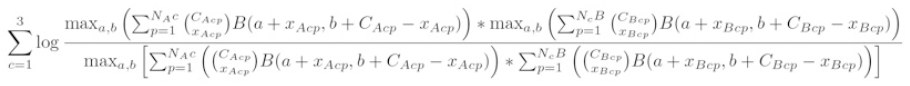

After correction for multiple testing using Storey’s method[56], an FDR threshold of 0.01 defined statistically different methylated regions (DMRs) between two strains.

Additionally, this method allowed calling differential methylation in a region for each context separately by computing *P* values as described above without summing over the contexts (*c* = 1, 2 or 3). We termed resulting DMRs CG-DMRs if the methylation at only CG sites within this region was statistically significantly different, and similarly CHG-DMRs and CHH-DMRs.

### Grouping differentially methylated strains in each region

For 13 strains there are at maximum 78 pairwise comparisons per region. To summarize pairwise comparisons and obtain epiallele frequencies, we assigned strains into differentially methylated groups. To achieve such clustering, we constructed a graph for each region where strains were represented as vertices and connected to other strains by an edge if the region was identified as a DMR between them (Figure S9b). We assume that strains within a group are then similarly methylated. The task is to find the smallest number of groups of vertices so that no two strains within a group are connected by an edge.

We set up a custom algorithm, which iteratively solves the “vertex coloring problem” for an increasing number of different colors, starting with two and quitting once all strains could be successfully assigned a color (Figure S9b). In each iteration, strains were processed in descendent order of their degree (i.e. number of edges it is connected to). Each strain was assigned to all possible colors that did not invoke a collision. Subsequently, the algorithm continued recursively to assign the color of the next strain.

Each strain had 3 context-dependent means of its beta binomial distributions per region (termed *strain means* from now on). We roughly approximated each group’s mean methylation values (*group means*) as the mean values of all *strain means* within a group. The *grouping diversity* describes the accumulated absolute differences between the *strain means* and their respective *group means* divided by the number of strains. As an example, consider Figure S9b. For simplicity, it only displays methylation rates for one out of three contexts. In the real data, the respective values were accumulated over all three contexts. The *group mean* for the blue strains in the example is (89+90+90+93+87)/5 = 89.8% and for the white strains 52%. The *grouping diversity* of the clustering shown here would be (from strains A to K): (|56-52| + |59-52| + |64-52| + |89-89.8| + |41-52| + |93-89.8| + |90-89.8| + |45-52| + |47-52| + |90-89.8| + |45-52| + |87-89.8|) / 11 = 2.84.

If there was more than one possible grouping of the strains, we chose the one with the lowest *grouping diversity*. A strain with no edges (i.e. which is not statistically differentially methylated to any other strain) was assigned into the group to which the accumulated absolute difference between its *strain mean* and the *group mean* was lowest. In the example of Figure S9b, strain L is grouped to the blue strains because its mean methylation value (81%) is closer to the blue *group mean* (90%) than to the white one (52%).

This procedure summarized the ∼221,000 DMRs of all pairwise strain comparisons into 11,323 DMRs between groups of strains.

### Testing regions for differential methylation between groups of strains

Once grouped, the same statistical test as for differential methylation between two strains was used to test groups of strains. Beta binomial distributions were approximated using the read counts of all strains in a group as if they were replicate data. This procedure identified 10,645 groups of regions showing significantly different methylation. Because the method used for the selection of the regions to perform the differential test can result in overlapping regions, DMRs can still overlap each other. From sets of overlapping DMRs, the non-overlapping DMR(s) with the lowest ‘grouping diversity’ was (were) retained, resulting in 4,821 final DMRs. For the vast majority of DMRs (98%), strains were classified into two groups, i.e. there are only two epialleles.

### Heritability analysis of methylated regions

For each differentially methylated region, we considered a linear mixed model to estimate the proportion of variance that is attributable to genetic effects (heritability) and its standard error. The approach is similar to variance component models used in GWAS, e.g. refs. [57,58]. Briefly, we considered the log average methylation rate of DMRs as phenotype and assessed the variance explained by genotype using a Kinship model constructed from all segregating genetic variants. We considered only DMRs and genetic polymorphisms that had no missing data in all accessions.

### Population structure analysis

We identified non-synonymous SNPs using SHOREmap_annotate [59] and excluded them from population structure analyses. We ran STRUCTURE v.2.3.4 [60] with *K*=2 to *K*=9 with a burn-in of 50,000 and 200,000 chains for 10 repetitions and determined the best *K* value using the Δ*K* method [61]. The phylogenetic network was generated using SplitsTree v.4.12.3 [62].

### Mapping to genomic elements

We used the TAIR10 annotation for genes, exons, introns and untranslated regions; transposon annotation was done according to [63]. Positions and regions were hierarchically assigned to annotated elements in the order CDS > intron > 5’ UTR > 3’ UTR > transposon > intergenic space. We defined as intergenic positions and regions those that were not annotated as either CDS, intron, UTR or transposon.

Positions were associated to the corresponding element when they were contained within the boundaries of that element. (D)MRs were associated to a class of element if they overlapped with that class of element; a (D)MR could only be associated to one class of element. When summing up basepairs of an element class covered by (D)MRs, the number of basepairs of a (D)MR overlapping with that class of element were considered. In that case the space covered by a (D)MR could be assigned to different classes of elements, while each basepair of the (D)MR could be assigned to only one class.

### Overlapping region analysis

We tested for significant overlap of DMRs using multovl version 1.2 (Campus Science Support Facilities GmbH (CSF), Vienna, Austria). We reduced the genome space to the basepair space covered by MRs identified in at least one HPG1 accession. DMRs were considered in the analysis if their start and end positions were contained within the MR space. DMRs that only partially overlapped with the MR space were trimmed to the overlapping part. Overlap between DMRs from different datasets was analyzed by running 100,000 permutations of both DMR sets within the MR basepair space. multovl commands are listed in Supplementary File 1.

### Processing and alignment of RNA-seq reads

Reads were processed in the same way as genomic reads, except that trimming was performed from both read ends. Filtered reads were then mapped to the TAIR9 version of the *Arabidopsis thaliana* (http://www.arabidopsis.org) genome using Tophat version 2.0.8 with Bowtie version 2.1.0 [64,65]. Coverage search and microexon search were activated. The command lines for Tophat are listed in Supplementary File 1.

### Gene expression analysis

or quantification of gene expression we used Cufflinks version 2.0.2[66]. We ran a Reference Annotation Based Transcript assembly (RABT) using the TAIR10 gene annotation (ftp://ftp.arabidopsis.org/home/tair/Genes/TAIR10_genome_release/TAIR10_gff3/) supplied with the most recent transposable element annotation [63] Fragment bias correction, multi-read correction and upper quartile normalization were enabled; transcripts of each sample were merged using Cuffmerge version 2.0.2, with RABT enabled. For detection of differential gene expression we ran Cuffdiff version 2.0.2 on the merged transcripts; FDR was set to < 0.05 and the minimum number of alignments per transcripts was 10. Fragment bias correction, multi-read correction and upper quartile normalization were enabled. The command lines for the Cufflinks pipeline are listed in Supplementary File 1. Analysis and graphical display of differential gene expression data was done using the cummeRbund package version 2.0.0 under R version 3.0.1.

### Data visualization

When not mentioned otherwise in the corresponding paragraph, graphical displays were generated using R version 3.0.1 (www.r-project.org). Circular display of genomic information in Figure 2a was rendered using Circos version 0.63 [67].

### Phenotyping

Leaf area was determined using the automated IPK LemnaTec System and the IAP analysis pipeline [68]. Plants were grown in a controlled-environment growth-chamber in an alpha-lattice design with eight replicates and three blocks per replicate, taking into account the structural constraints of the LemnaTec system. Each block consisted of eight carriers, each carrying six plants of one line. Stratification for 2 days at 6°C was followed by cultivation at 20/18°C, 60/75% relative humidity in a 16/8 h day/night cycle. Plants were watered and imaged daily until 21 days after sowing (DAS). Adjusted means were calculated using REML in Genstat 14^th^ Edition, with genotype and time of germination as fixed effects, and replicate|block as random effects.

### Data accessibility

The DNA and RNA sequencing data have been deposited at the European Nucleotide Archive under accession number XXX and XXX. A GBrowse instance for DNA methylation and transcriptome data is available at (to be released upon publication). DNA methylation data and MR coordinates have also been uploaded to the EPIC-CoGe browser (data will be made publicly available upon publication of the manuscript).

## ACKNOWLEDGEMENTS

We are grateful to C. Lanz for help with the Illumina sequencing, Q. Song and A. Smith for making the source code of the Hidden Markov Model implementation available, the group of V. Colot for sharing the Col-0 MeDIP-Seq data, and C. Klukas for assistance with the processing of the phenotyping data. We thank R. Schwab for critical reading of the manuscript. This work was supported by a Marie Curie FP7 fellowship (O.S.), grant NIH GM083068 (J.B.), FP7 Collaborative Project AENEAS (contract KBBE-2009-226477), a Gottfried Wilhelm Leibniz Award of the DFG, and the Max Planck Society (D.W.).

## AUTHOR CONTRIBUTIONS

C.B., J.H., T.A., J.B., and D.W. conceived the study; C.B. and R.C.M. performed the experiments; J.H., C.B., J.M., O.S., K.S. analyzed the data; J.F. implemented the data visualization; K.B. provided advice on statistical analysis; and C.B., J.H. and D.W. wrote the paper with contributions from all authors.

## AUTHOR INFORMATION

Correspondence and requests for materials should be addressed to D.W.

## SUPPORTING INFORMATION

Supplementary information is linked to the paper.

## Supplementary Online Material

### Century-scale methylome stability in a recently diverged *Arabidopsis thaliana* lineage

*Jörg Hagmann, Claude Becker, Jonas Müller, Oliver Stegle, Rhonda C. Meyer, Korbinian Schneeberger, Joffrey Fitz, Thomas Altmann, Joy Bergelson, Karsten Borgwardt, Detlef Weigel*

### Genome analysis of HPG1 individuals

To answer how heritable epigenetic differences are affected by long-term exposure to fluctuating and diverse environmental conditions, we wished to identify a nearly isogenic lineage among natural accessions of *A. thaliana* that had diverged for at least several dozens of generations. In the natural range of the species in Eurasia, nearly isogenic individuals are generally found only at single sites, many of which are transient [1,2]. An exception is North America, where about half of all individuals sampled appear to be identical when genotyped at 139 genome-wide markers [2], in agreement with a limited number of founders that were introduced from Europe in historic times [3]. We here refer to this lineage as haplogroup-1 (HPG1), because it dominates the North American population.

We selected 13 HPG1 individuals from seven locations in the Eastern Lake Michigan area, from one location in Western Illinois, and one location on Long Island; the median distance between sites was 155 km. The set consisted of pairs of accessions from each of four sites, and single individuals from the other five sites (Figure 1a, Table S1).

We assessed genetic divergence among the 13 HPG1 lines by Illumina paired-end sequencing (see table below).

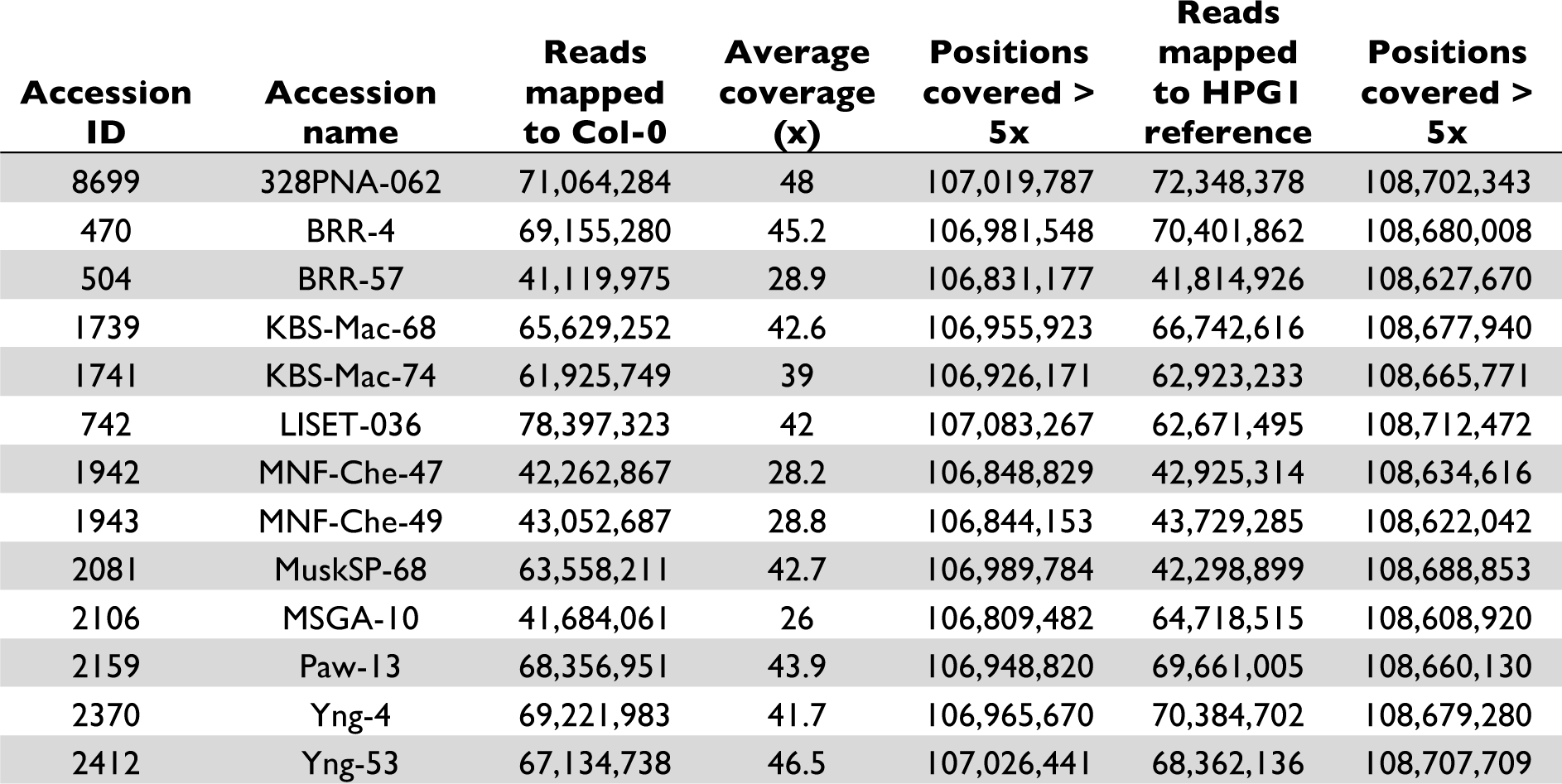

We identified an initial set of common single-nucleotide polymorphisms (SNPs), small-scale indels and structural variants (collectively referred to as SVs henceforth) by read alignment to the Col-0 reference genome. We iteratively built an HPG1 pseudo-reference genome through integration of the common variants into the Col-0 genome, re-alignment of HPG1 reads to this new reference and re-calling of SNPs and SVs (Figure S1a, c), following the rationale of Gan and colleagues [4]. After two iterations, the number of common variants had increased by 12% and the number of reads that could not be mapped had decreased by a third (Figure S1b). We ultimately identified 670,979 common SNPs and 170,998 SVs compared to the Col-0 reference (Table S2). Considering the corresponding nucleotides in the close relative *A. lyrata* as the ancestral states [5], a little bit more than half of the SNPs at positions alignable to the *A. lyrata* genome were classified as derived in the HPG1 population, close to what would be expected for a comparison of an arbitrary set of accessions (i.e., the HPG1 ancestor and Col-0) (Table S2).

We additionally called variants in each of the 13 individuals based on the HPG1 pseudo-reference genome. Compared to the common variants, a much smaller number, 1,354 SNPs and 521 SVs, segregated in the HPG1 population (Table S3), confirming that the 13 strains were indeed closely related. As for common variants, segregating variants were fairly equally distributed across all ten chromosome arms (Figure S2). Eighteen percent of both segregating and common SNPs mapped to genes, and 30% of segregating and 35% of common SNPs to non-transposable element intergenic regions (Figure S3; Table S3). This was similar to SNPs in other natural accessions [6], indicating that HPG1 is representative of natural accessions of *A. thaliana*. On average, two HPG1 accessions differed by 294 SNPs.

### Primary analysis of methylation

We performed whole methylome bisulfite sequencing to an average depth of 18x per strand (Table S4) on two pools consisting of 8-10 individuals per accession. The pooling was performed to reduce variation in methylation rate caused by stochastic fluctuations in coverage or read sampling biases. Using the HPG1 pseudo reference genome instead of the Col-0 reference genome increased the number of cytosines sufficiently covered for statistical analysis by 5% on average, and the number of positions called as methylated by 7% (Table S4). Of 24 million cytosines that were covered by at least three independent reads, on average 2.5 million were methylated per strain (Table S4).

For the analysis of differentially methylated positions (DMPs), we considered only cytosines that were covered by at least three independent reads in at least seven strains, 21.1 million in total. We found mostly DMPs in the CG context (97%). This can be partially explained by our statistical test, which more easily identifies large differences in methylation, as is typical for variation at CG sites (see discussion in SOM of ref. [7]). In addition, stable silencing of repeats and TEs that are rich in CHG and CHH sites may lead to a similar patterns. To assess epiallele frequencies, we compared methylation in each of the 12 accessions from near Lake Michigan to the geographic outlier LISET-036 from Long Island, which we selected as a reference strain. Sixty-one percent of CG-DMPs were recurrent in at least two independent Lake Michigan accessions (Figure S6a), which is almost double of what we had previously observed for ten equidistant greenhouse-grown MA lines [7]. This can be partly explained by the fact that we sequenced four pairs of strains from the same location. Forty-five percent of all CG-DMPs with a frequency of 2 were attributable to such pairs, while 6% would be expected if they were randomly distributed across strains.

### Estimating DMP accumulation rates

We had previously sequenced the genomes of only five of the 12 MA lines for which we had reported DMPs [7], [8]. We therefore generated additional genome sequence data for all ten MA lines in generation 31, counting from the founder plant of the population, as well as from the two lines in generation 3 [7]. To increase the number of data points in the low range of genetic differences, we inferred the number of SNPs between siblings (which had been included in the previous MA methylome analyses [7]) from the mutation rates determined by Ossowski and colleagues on the same lines. The greater variance in DMP rates and the more rapid initial increase in DMPs in the MA lines in Figure 2b is presumed to be due to the methylome data having come from individual plants, instead of from pools of individuals as for the HPG1 lines. By pooling strains, low frequency epimutations are diluted and less likely to be detected. This assumption is corroborated by the fact that we see only 46 DMPs between replicates of HPG1 pools compared to about 1,300 in replicates of the MA lines. To further investigate this, we compared the number of DMPs after *in silico* pooling of individuals. We first combined data from two siblings of MA line 30-39 or 30-49 in generation 31 with two siblings of their generation 32 offspring. We then calculated DMPs in comparison of pooled data from two times two individuals from two different generation 31 MA lines. We compared the results with those from individual comparisons of all 16 pairwise comparisons between the four early- and four late-generation individuals. The number of DMPs distinguishing pools was at the lower end of the DMP distribution from the individual comparisons (Figure S7). Hence, it is likely that we underestimate the epimutation rate of the HPG1 accessions (Figure 2b).

Moreover, we assumed the same genetic mutation rate in the two populations. A potentially faster genetic mutation rate in the wild would result in a steeper slope of the HPG1 curve, if plotted against number of generations. Finally, the initial increase of the HPG1 epimutation rate is based on only few comparisons between strains from the same sampling site, which might not be sufficient for an accurate estimate.

### Methylated regions

The value of an approach that defines methylated regions (MRs) before identifying differentially methylated regions (DMRs) has been demonstrated before with a Hidden Markov Model (HMM) method developed for the analysis of methylated-DNA-immunoprecipitation followed by array hybridization (MeDIP-chip) [9]. We therefore implemented a two-state HMM that allows MR identification based solely on within-genome variation in methylation rate. The model, based on one for animal DNA methylation data [10], learns methylation rate distributions for both an unmethylated and a methylated state for each sequence context separately (CG, CHG and CHH) while simultaneously estimating transition probabilities between the two states from genome-wide data. On the trained model, the most probable path of the HMM along the genome is then used to define regions of high and low methylation. Applied to the HPG1 population, the HMM identified on average 32,530 (SD = 1,629) MRs in each strain. The unified MRs had a median length of 122 bp (mean = 649 bp), with a maximum length of 87.7 kb (Figure S8a; Table S5).

For validation we compared data generated from Col-0 (see below) to data from methylated-DNA immunoprecipitation followed by sequencing (MeDIP-seq; Vincent Colot and co-workers, pers. communication). Of the genome space enriched in MeDIP-seq, 89% was classified as MR by our HMM approach.

To also evaluate whether the identified MRs sufficiently capture methylated sites within gene bodies consisting almost exclusively of CG sites, we tested how many MRs fall into gene body methylated genes as defined previously [11]. We re-implemented their method and called between 4,330 and 4,626 gene body methylated (BM) genes and between 14,998 and 15,489 unmethylated (UM) genes for the HPG1 strains. These figures are similar to the 4,361 BM and 15,753 UM genes reported in ref. [11] for Col-0. A quarter of the BM genes identified in this study and in ref. [11] did not overlap, which may be due to genetic differences and/or different sequencing depths and analysis pipelines. MRs in this work overlap with 58% of the HPG1 BM genes. This compares with an overlap of MeDIP domains of Col-0 (Colot lab, see above) of only 42% with Col-0 BM genes. Moreover, the concepts of our approach and that in ref. [11] differ considerably: by modeling the density of methylated sites within a gene with a binomial distribution allowing only little variance, genes with slightly increased density of methylated sites compared to the global average are quickly classified as BM in the method of ref. [11]. Such sites can still be located far apart from each other. In contrast, MRs are called by our HMM-based method only when there is a locally restricted, dense region of methylated sites. BM genes that are covered by MRs have a higher density of methylated sites compared to BM genes without overlapping MRs (Figure S8d).

### Differentially methylated regions

To identify DMRs, we performed pairwise comparisons of overlapping MRs or parts thereof that were classified either (i) as in high methylation state in both accessions of a pair, or (ii) as high methylation state fragments in one and low methylation state in the other accession. For each DMR we then assigned all accessions to groups, based on their significant methylation differences (Figure S9). We expected to find fewer DMRs in regions that had a high methylation state in both tested accessions. In agreement, only 0.4% of those tested fragments (31,531 out of 7,355,716) were significantly differentially methylated. In contrast, we identified as differentially methylated 4.4% (107,988 out of 2,450,278) of fragments where the tested accessions had been assigned to contrasting methylation states. Consolidation of all pairwise comparisons, resolving overlapping DMRs and differentiality testing after grouping resulted in a final set of 4,821 DMRs with an average length of 159 bp (Figure S8a; Figure S9; Table S6).

Our sensitive statistical test classified as differential some regions with low variance and only subtle methylation difference; we therefore defined as highly differentially methylated regions (hDMRs) with potentially greater biological relevance all DMRs that were longer than 50 bp and that showed a more-than-three-fold difference in methylation rate in at least one sequence context, when considering at least five cytosines of that context (Figure S9). In addition, the overall methylation rate of the DMR in the more highly methylated strain had to be greater than 20%. Of 3,909 size-filtered DMRs, 3,199 (80%) were classified as hDMRs (Table S7). The grouping of hDMRs yielded similar epiallele frequencies as for the DMPs (54% with frequency larger than 1; Figure 2f). The independent analysis of cytosines from different contexts allowed us to call context-specific (h)DMRs. In 71% of DMRs and 76% of hDMRs, only cytosines in the CG context significantly differed in their methylation rate. Only a minority, 15% of DMRs and 7% of hDMRs, showed highly variable methylation in more than one cytosine context (Figure 2g).

In contrast to previously used methods for the analysis of whole-genome bisulfite sequencing data from plants, our HMM for the detection of MRs does not require information about methylation differences at the single-site level. By first identifying blocks of methylation, our approach limits the number of tested regions to the methylated space of the genome, thereby reducing the multiple testing problem and the requirement for arbitrary filters. Importantly, limiting DMR detection to HMM-identified MRs revealed that the location of DMRs in the genome follows the overall distribution of methylation. Most of these DMRs thus overlap with TEs and intergenic regions, which is in contrast to previously published DMRs relying on user-defined criteria including arbitrary sliding windows or DMPs [7,12–21]. The HMM analysis also revealed that non-CG methylation is almost exclusively organized in regions of contiguous DNA methylation. We suggest such an approach to identifying MRs be applied to bisulfite sequencing data in future studies.

### Analysis of LISET-036 specific hDMRs

Strain LISET-036 was the most different when strains were clustered by CHG-DMPs, CHH-DMPs and DMRs. Since CHG and CHH-DMPs constituted only a minor fraction of all DMPs (∼3%), we focused on hDMRs private to the HPG1 strains to investigate the possible basis of LISET-036 being an outlier. While LISET-036 had the most private hDMRs among all accessions (Figure S18a), their spectrum in terms of context and overlap with genomic features did not deviate from that of the other strains (Figure S18b). 44 LISET-036 private hDMRs overlapped with genes and 30 overlapped with the gene-adjacent regions, defined as 1,000 bp upstream or downstream of genes. The only GO term for which these 74 genes were enriched was “intrinsic to membrane” (p-value 0.01). However, there were no overlapping differentially expressed genes. Taken together, there is little evidence for a pronounced phenotypic effect of the LISET-036-specific epivariants.

### Differential gene expression and epigenetic variation

We performed RNA-seq (Table S8) on rosette leaves of all 13 strains and identified 251 differentially expressed (DE) genes across all possible pairwise comparisons (Table S9). A majority of these genes were identified as DE in more than one comparison. Gene Ontology (GO) analysis identified several defense-related GO terms to be over-represented (p << 0.001); which may be linked to these genes evolving fast [22]. As could be expected from the small numbers of DE genes, clustering of accessions based on expression of all genes revealed no structure reflecting genetic distance or geographical origin. When we limited the clustering to DE genes, however, accessions originating from the same geographical location, except Yng-4 and Yng-53, clustered together (Figure S11a). A similar observation could be made when counting the number of DE genes per comparison: while accessions from the same site generally showed no or only few DE genes, comparison of accessions from different sites revealed up to 149 differences. The two Yng strains accounted for most of the DE genes identified in pairwise comparisons (Figure S11b). Although we identified some loci where changes in contiguous stretches of DNA methylation correlated with alterations in transcriptional activity (Table S10), we did not observe a general relationship between these two features, suggesting either that transcriptional differences in the HPG1 population is mostly due to DNA mutations, or due to epigenetic changes independent of DNA methylation.

### Similarity of epigenetic variation profiles in independent populations

We used the data we had previously generated on the greenhouse-grown MA lines [7] to investigate whether the MA epigenetic variants were similar to the HPG1 variants. Almost half of all positions that had been classified as DMPs in the MA lines and that were sufficiently covered in the HPG1 accessions were also polymorphic in the HPG1 accessions (41%; p<<1 x 10^−5^, Fisher’s exact test). Similarly, about a third of HPG1 DMPs were also MA DMPs (29%; p<<1 x 10^−5^) (Figure 3a).

We recalled DMRs in the MA strains [7] using the methods implemented for the HPG1 population. We detected on average 22,868 MRs (SD = 1,282) per MA line, covering 22.3 Mb. Of 3,837 DMRs in the MA lines, 2,523 coincided with MRs detected in the HPG1 population. DMRs in the HPG1 population were 4-fold more likely to coincide with DMRs in the MA lines than with a random MR from this set (Figure 3b).

We also identified DMRs between one of the MA lines (30-39) and one of the HPG1 lines (MuskSP-68). These DMRs were also enriched in each of the two sets of within-population DMRs (MA or HPG1) (Figure 3b).

Finally, we performed a similar analysis with DMRs from 140 natural accessions that represent the entire range of the species [14]. While the overlap between those 53,752 DMRs and the HPG1 DMRs was greater than expected by chance (Z-score = 19.8; 100,000 permutations), most of the DMRs in the 140 accessions (9,994) do not even overlap with MRs in the HPG1 or MA population (Figure 3d). While this is almost certainly due at least in part to the very different DMR detection methods, it could also indicate that differential methylation in the global population was influenced by genetic variation to a larger degree than in the HPG1 accessions.

### Supplementary figure legends

**Figure S1.**
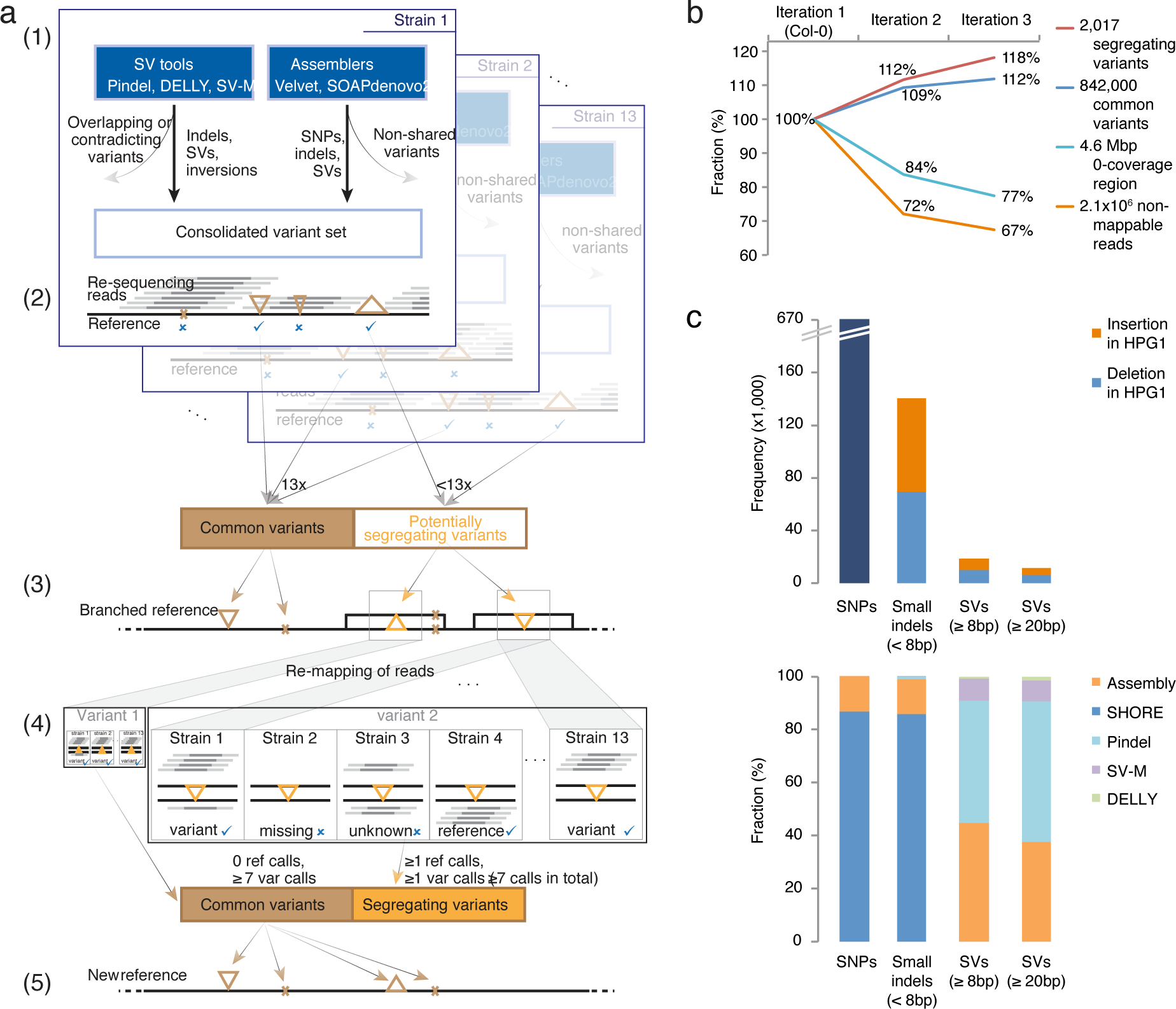
Iterative re-alignment strategy and statistics. (a) Iterative re-alignment approach to evaluate predicted variants and to build a HPG1 pseudo-reference genome. (1) For each strain, variants called by diverse structural variant detection tools and a local *de novo* assembly pipeline were combined into a consolidated variant set. (2) Variants with core read coverage were filtered out, the remaining variants were classified into common and potentially segregating. Brown triangles symbolize insertions/deletions, the brown X represents a SNP. (3) All common variants were incorporated into the reference genome. All segregating variants were introduced in branches of the reference genome, which incorporated polymorphisms linked by less than 107 bp. (4) After mapping the reads against the branched reference, a binomial test was performed in each strain and for each variable site to call the allele, i.e., to determine whether there was statistical evidence for the presence of only one haplotype covering the variant’s coordinates. Variants with the same non-reference allele call in all strains were considered as “common”; those with a reference call in at least one strain and a variant call in at least one other strain were classified as “segregating”. (5) All common variants from the previous step were incorporated into the new reference sequence, and a new iteration was started over from (1), or this new genome served as the HPG1 pseudo-reference genome after iteration 2. (b) Increase of detected variants and decrease of unsequenced genome space and unmappable reads by iterative read mapping. The legend on the right side denotes absolute values after iteration 2. The reference value (100%) derives from the mapping against the Columbia-0 genome sequence (TAIR9), and for common variants it is the number of variants leading to the genome of iteration 1. Thus, ∼842,000 common variants led to the genome of iteration 2, ∼864,000 to the genome of iteration 3. (c) Composition of common polymorphisms by variant type (top) and by detection tool (bottom). Variants found by more than one tool contributed to the count for all respective tools.

**Figure S2.**
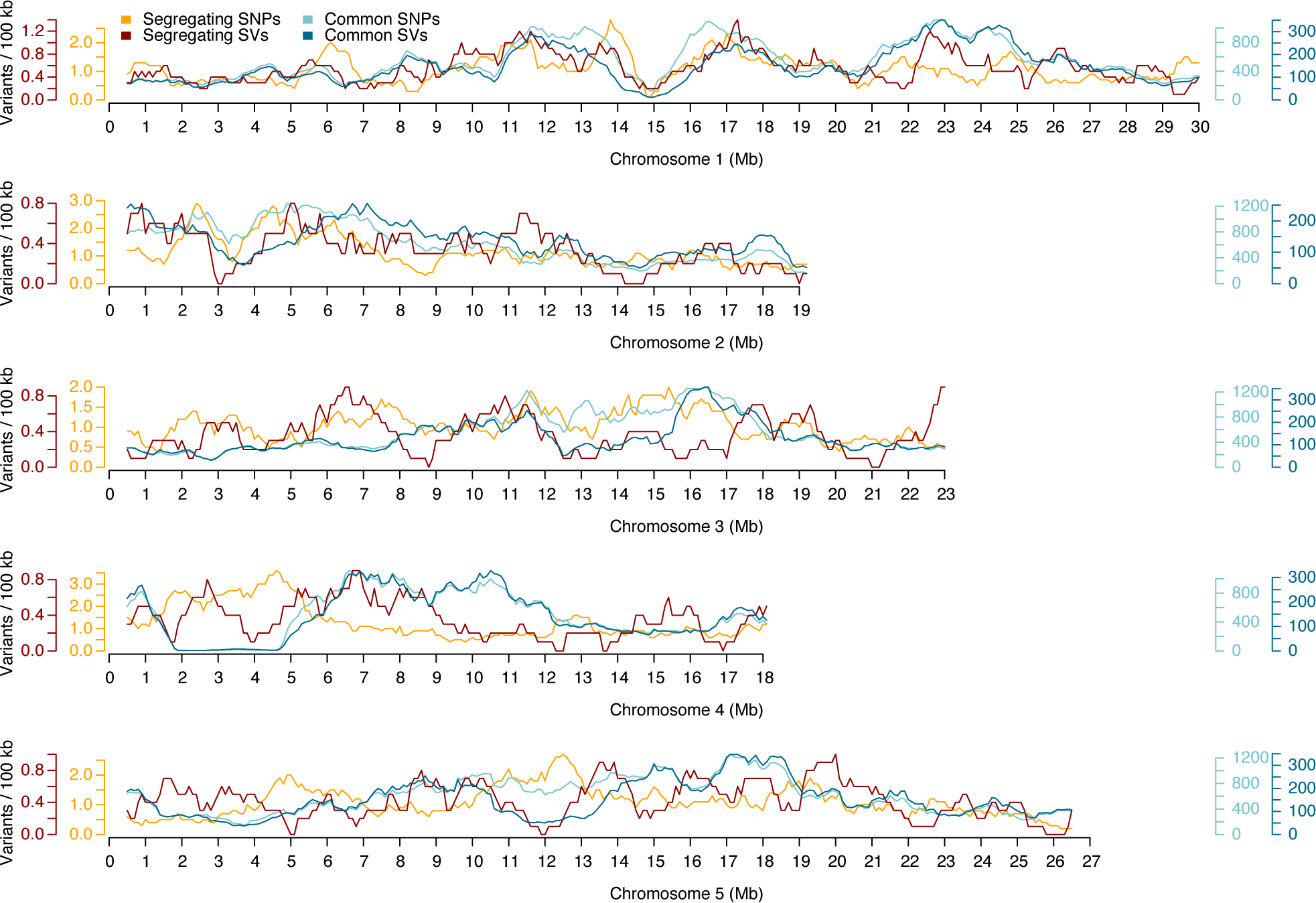
Distribution of genetic variants along the five chromosomes. Relative density of common variants in 100 kb sliding windows with a step size of 10 kb.

**Figure S3.**
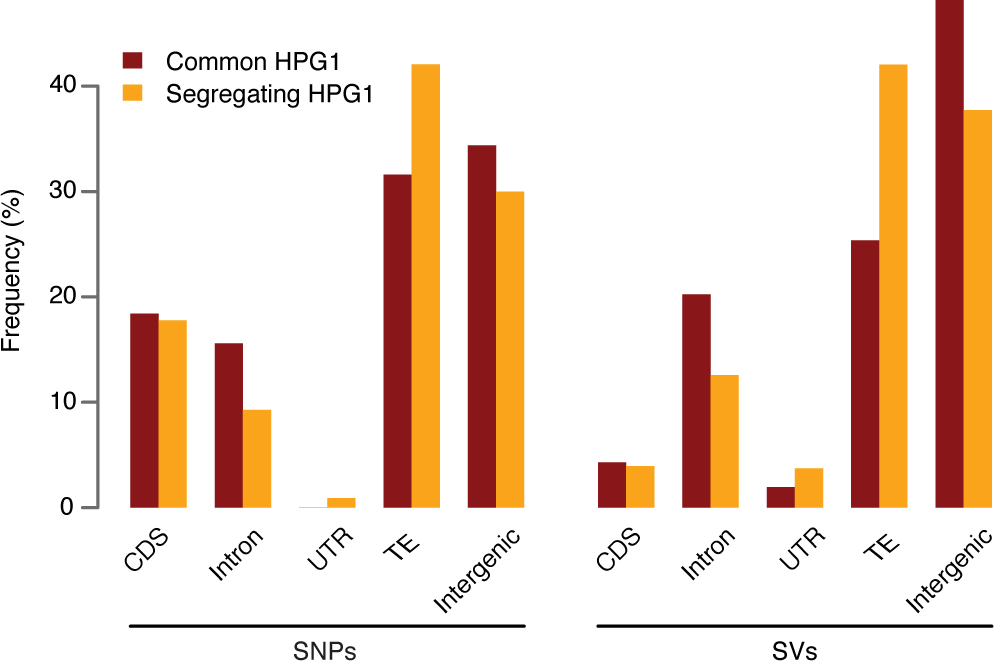
Annotation of genetic variants. Polymorphisms were hierarchically assigned to CDS > intron > 5’ UTR > 3’ UTR > transposon > intergenic.

**Figure S4.**
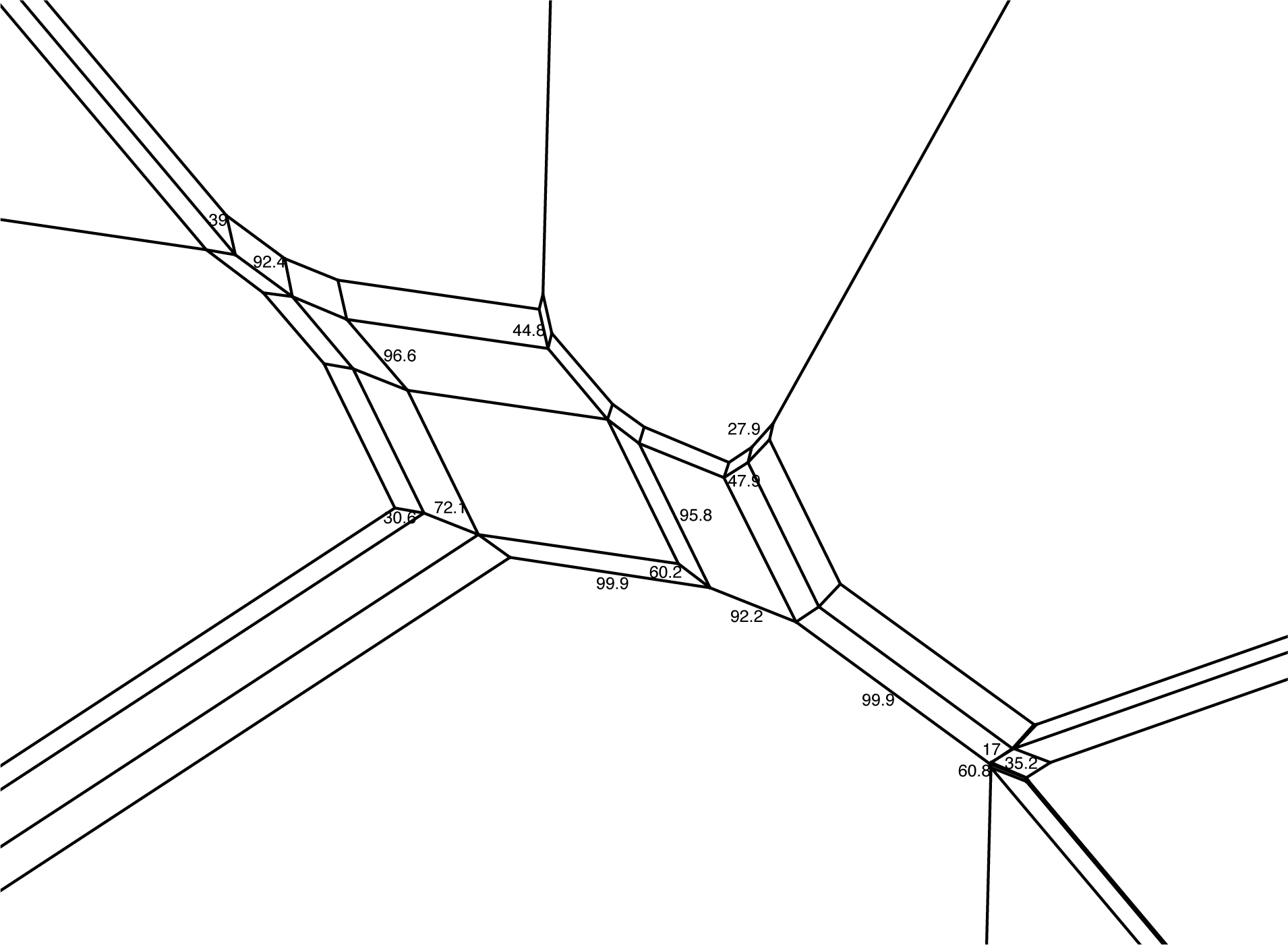
Magnification of the central area of the phylogenetic network in Figure 1c. Numbers indicate bootstrap confidence values (10,000 iterations).

**Figure S5.**
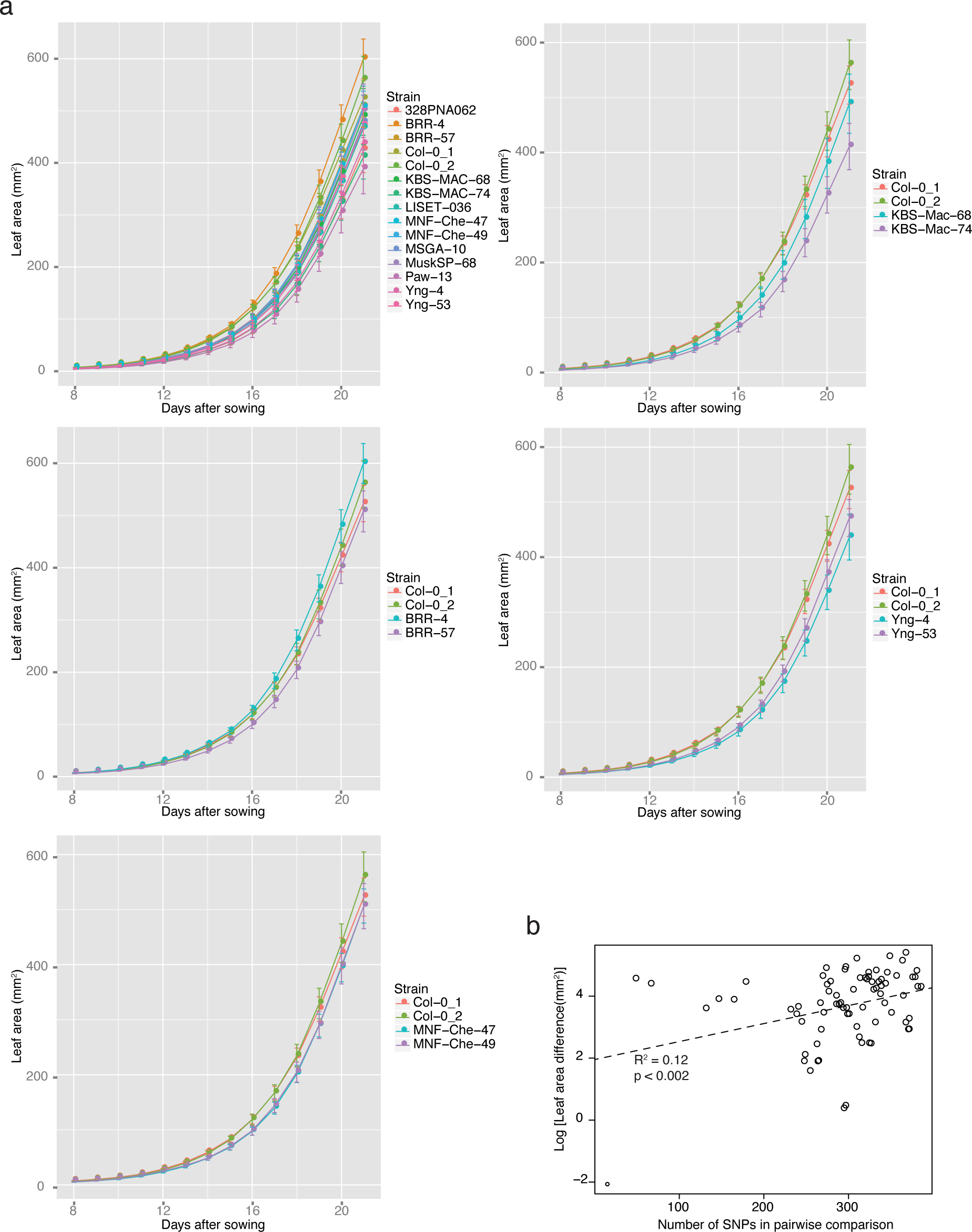
Phenotypic analysis. (a) Leaf growth measured over time (Materials and Methods). Error bars represent 95% confidence intervals. On average, 36 plants were measured per accession. (b) Correlation of genetic distance, represented by number of SNPs per pairwise comparison, and difference in leaf area at 21 days after germination.

**Figure S6.**
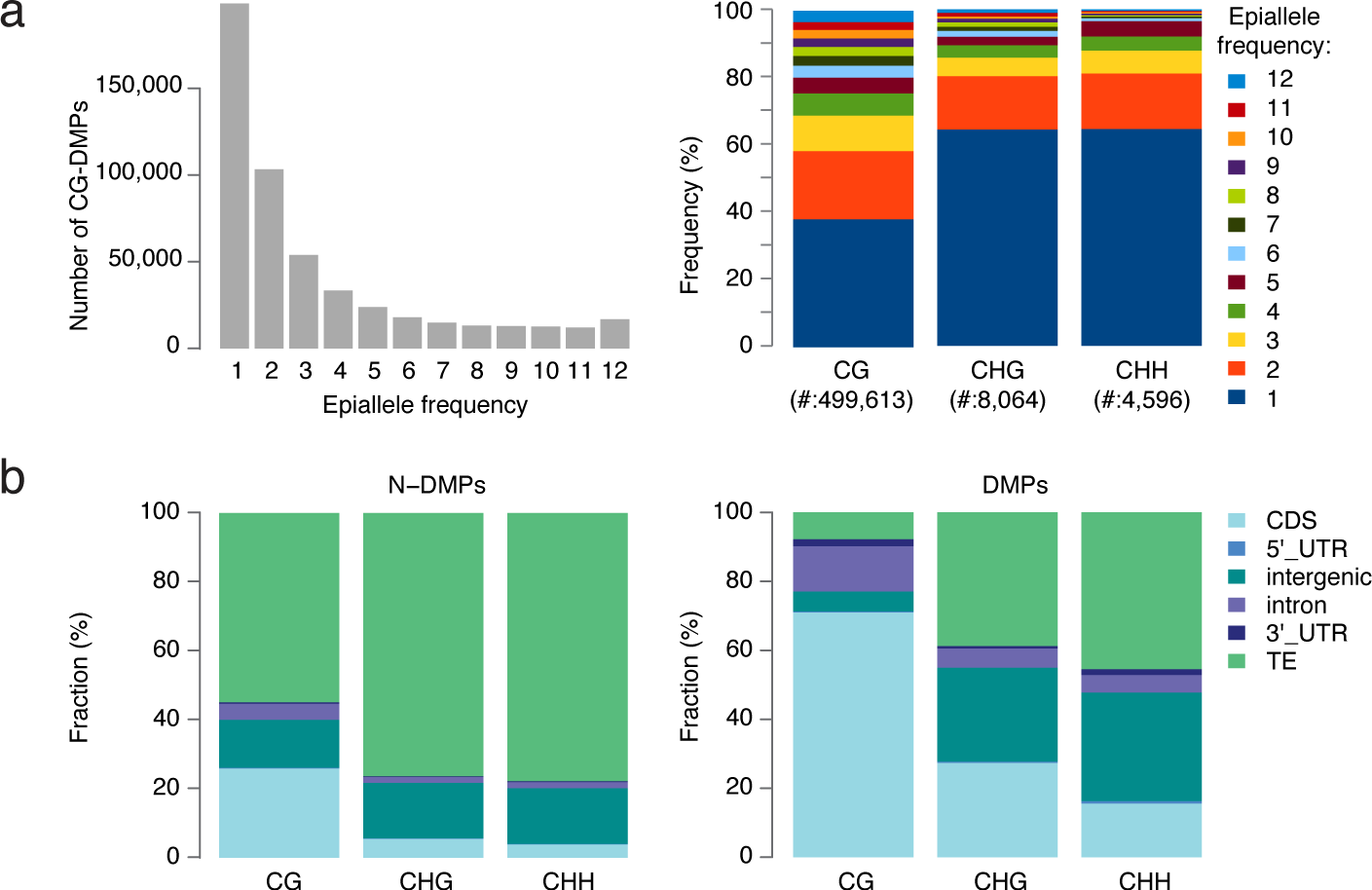
DMPs. (a) Epiallele frequencies of DMPs for CG sites only (left), and comparison of all three sequence contexts (right). (b) Annotation of ^5m^Cs and DMPs. Cytosines were hierarchically assigned to CDS > intron > 5’ UTR > 3’ UTR > transposon > intergenic.

**Figure S7.**
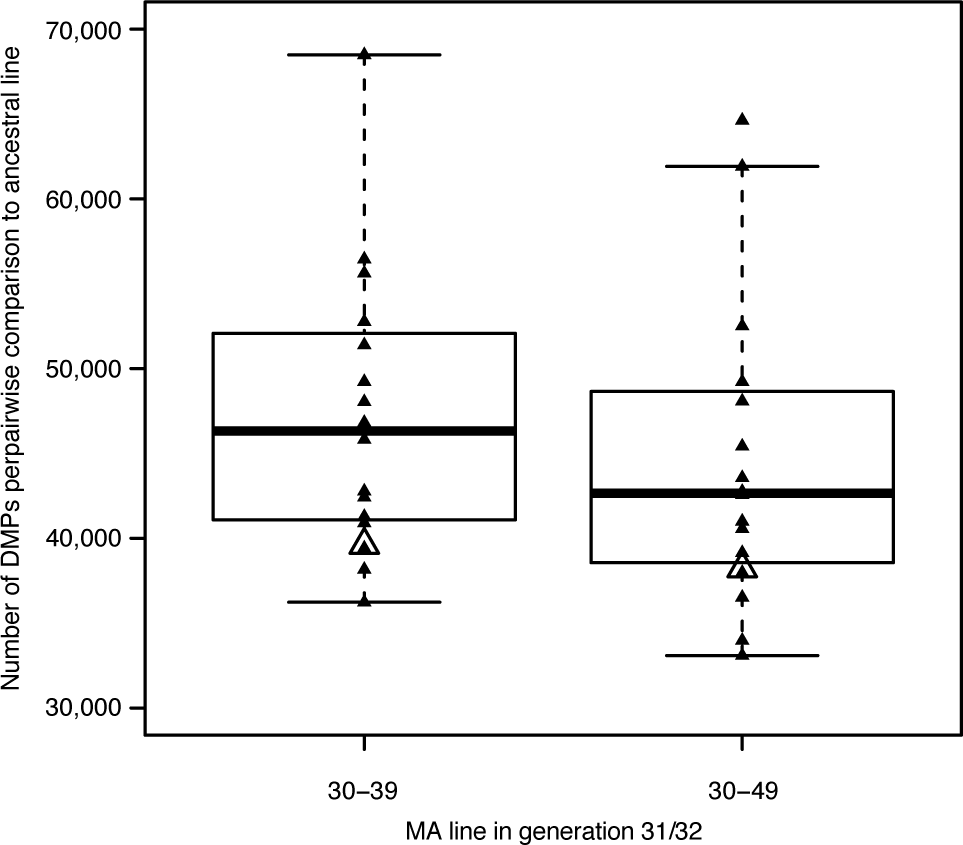
Effect of sample pooling on the number of identified DMPs. Small filled triangles and box-and-whisker plot indicate distribution of DMPs that were called by comparing individual plants of generations 31 and 32 of lines MA30-39 and MA30-49 with individual plants of MA0-4-26 and MA0-8-87, which represent generation 3 of two independent lines (16 comparisons in each group). The large unfilled triangles indicate the number of DMPs that were called when data from four lines of generations 31 and 32 were pooled in silico and compared against all four lines from generation 3. On average, a substantially lower number of DMPs is called with pooled data.

**Figure S8.**
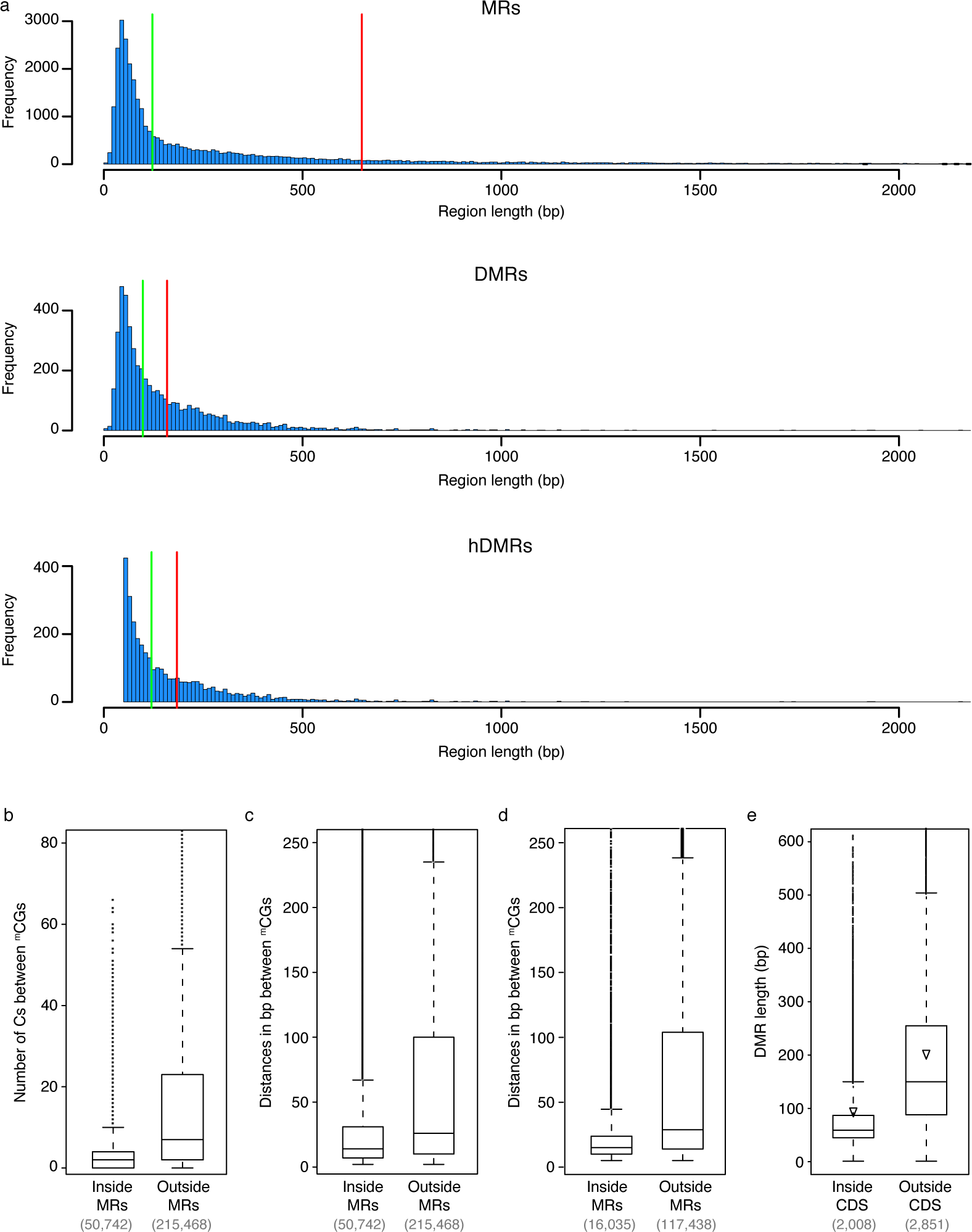
Characteristics of (differentially) methylated regions. (a) Histogram of the length of unified MRs, DMRs and hDMRs. The red and green lines indicate mean and median length, respectively. For better visibility, regions larger than 2,000 bp were excluded from the representation. The minimum size of hDMRs was 50 bp. (b)-(d) are based on data of one HPG1 strain (LISET-036). (b) Number of unmethylated cytosines in-between methylated CG sites within genes in dependence of whether these sequences are inside or outside of MRs. (c) Distances in bp between methylated CG sites within genes in dependence of whether these sequences are inside and outside MRs (minimal distance 2 bp). (d) Distances between methylated CG sites within body methylated (BM) genes identified with the method from ref. [11] (SOM: Methylated regions) and within genes not identified as BM (minimal distance 2 bp). (e) Length distributions of DMRs that overlap and that do not overlap coding regions. Triangles show mean values.

**Figure S9.**
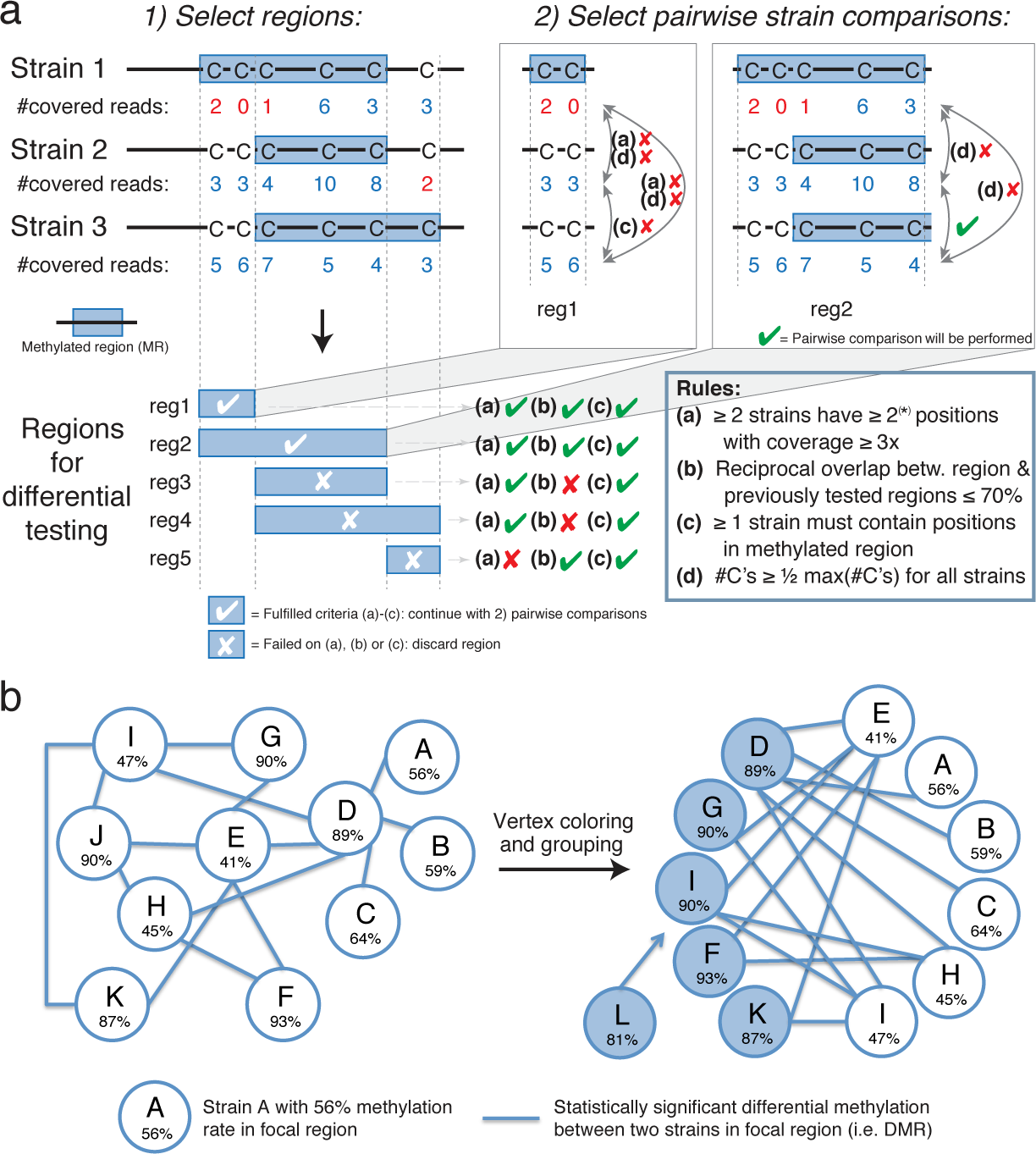
Selecting subregions of HMM-detected MRs to test for differential methylation. (a) Example illustrating the selection procedure of regions and pairwise strain comparisons to be tested for differential methylation. (*) For simplicity, the illustration uses a minimum number of covered sites of two reads per region (10 reads for the real data set). (1) We selected all possible regions where two strains presented different states of methylation (*reg1* to *reg5*) and applied filter criteria (a), (b) and (c). (2) If a region passed filters (a), (b) and (c) (in the example only *reg1* and *reg2*), criteria (a), (c) and (d) were checked for each pairwise comparison between strains on that region. Note, the selection of a region in (1) must not necessarily lead to a differential test between any two strains (e.g., *reg1*). Refer to the Material and Methods section for elaborate descriptions of criteria. (b) Assignment of strains to different groups based on differential methylation. Left: An exemplary DMR represented as a graph: strains are represented as nodes, edges reflect a statistically significant test between two strains. Right: Finding the minimal number of sets, where no edge connects nodes from two different groups is known as the colouring vertex problem. In this example, the solution is two sets of strains (blue and white nodes). Strains without statistically significant tests (e.g., strain L) are grouped into the set of strains where the difference between the strain’s and the group’s mean methylation rate is minimal.

**Figure S10.**
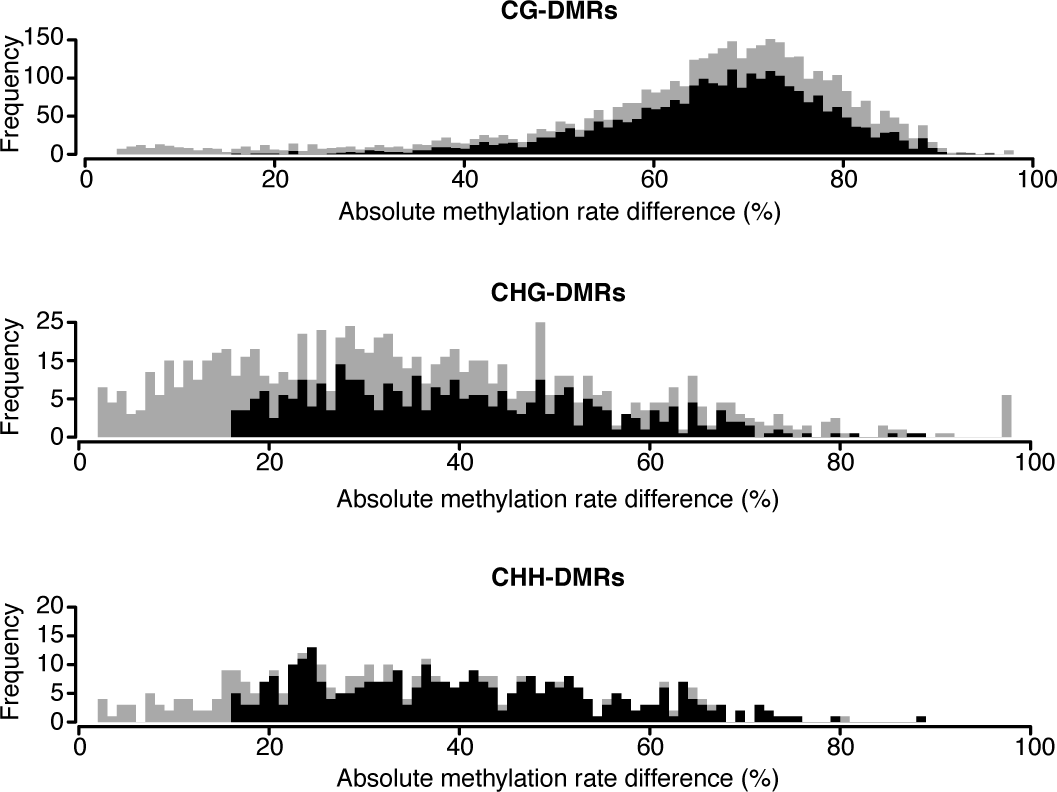
Difference in local methylation rate between lines classified as “low methylation” versus lines classified as “high methylation” over a given (h)DMR. Histograms of the absolute mean methylation rate differences of DMRs (grey) and hDMRs (black) of different sequence contexts.

**Figure S11.**
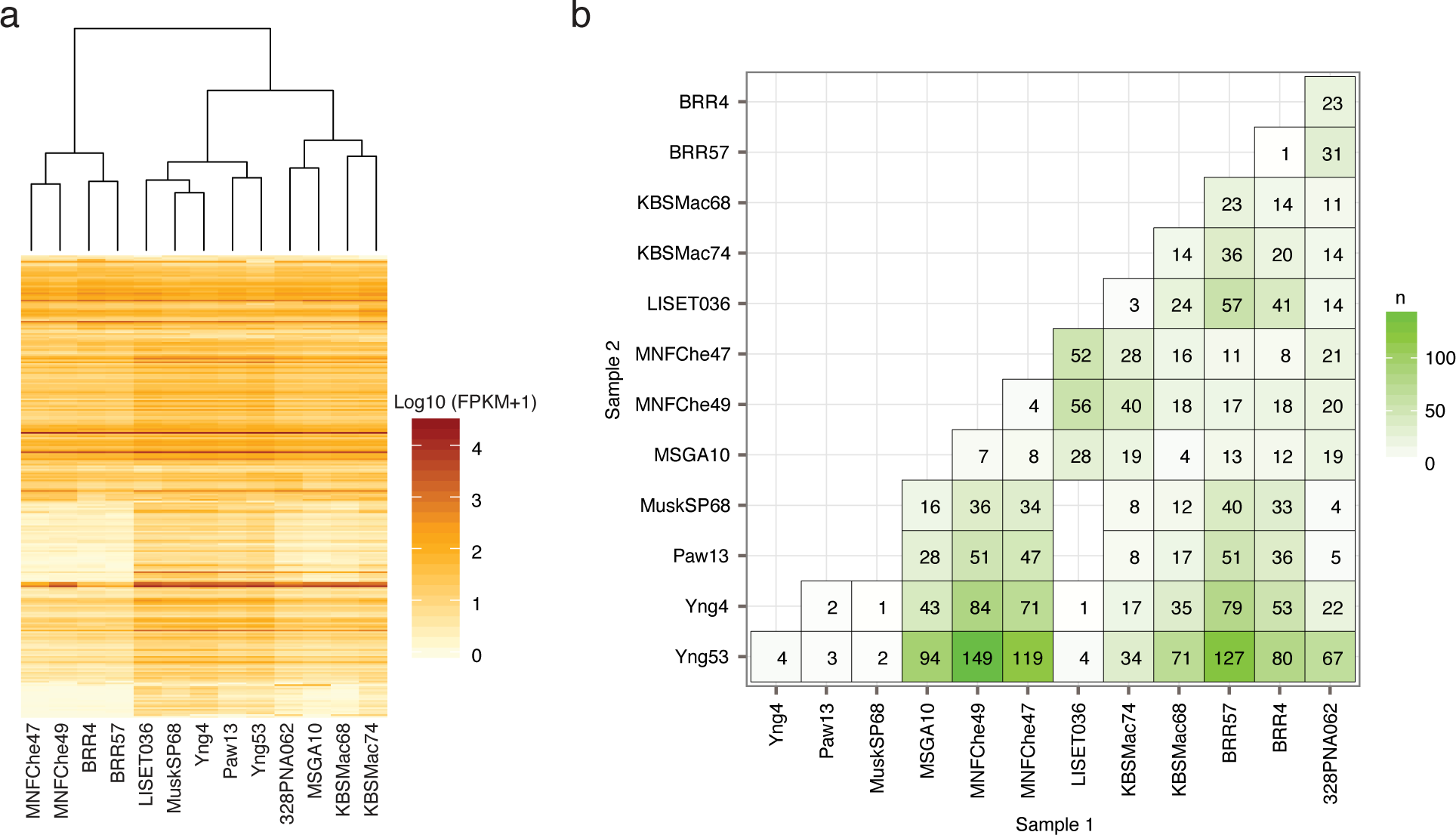
Differential gene expression. (a) Hierarchical clustering of HPG1 accessions by expression of differentially expressed genes. (b) Differentially expressed genes per pairwise comparison.

**Figure S12.**
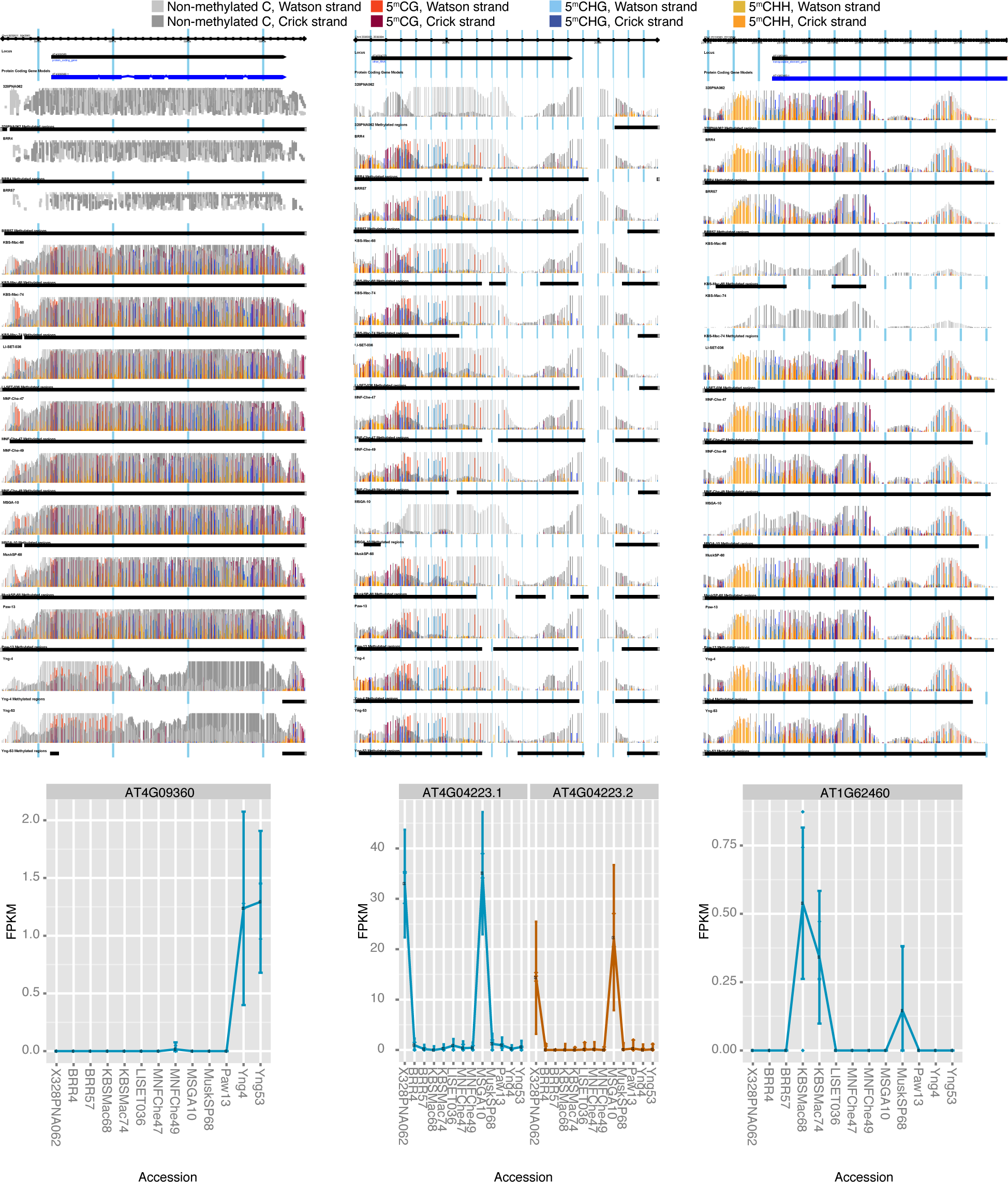
DMRs and gene expression. Examples of DMRs (top panel) overlapping with a protein-coding gene (AT4G09360, left), a non-coding RNA (AT4G04223, middle) and a transposable element (AT1G62460, right). The expression of the corresponding locus is represented in the bottom panel.

**Figure S13.**
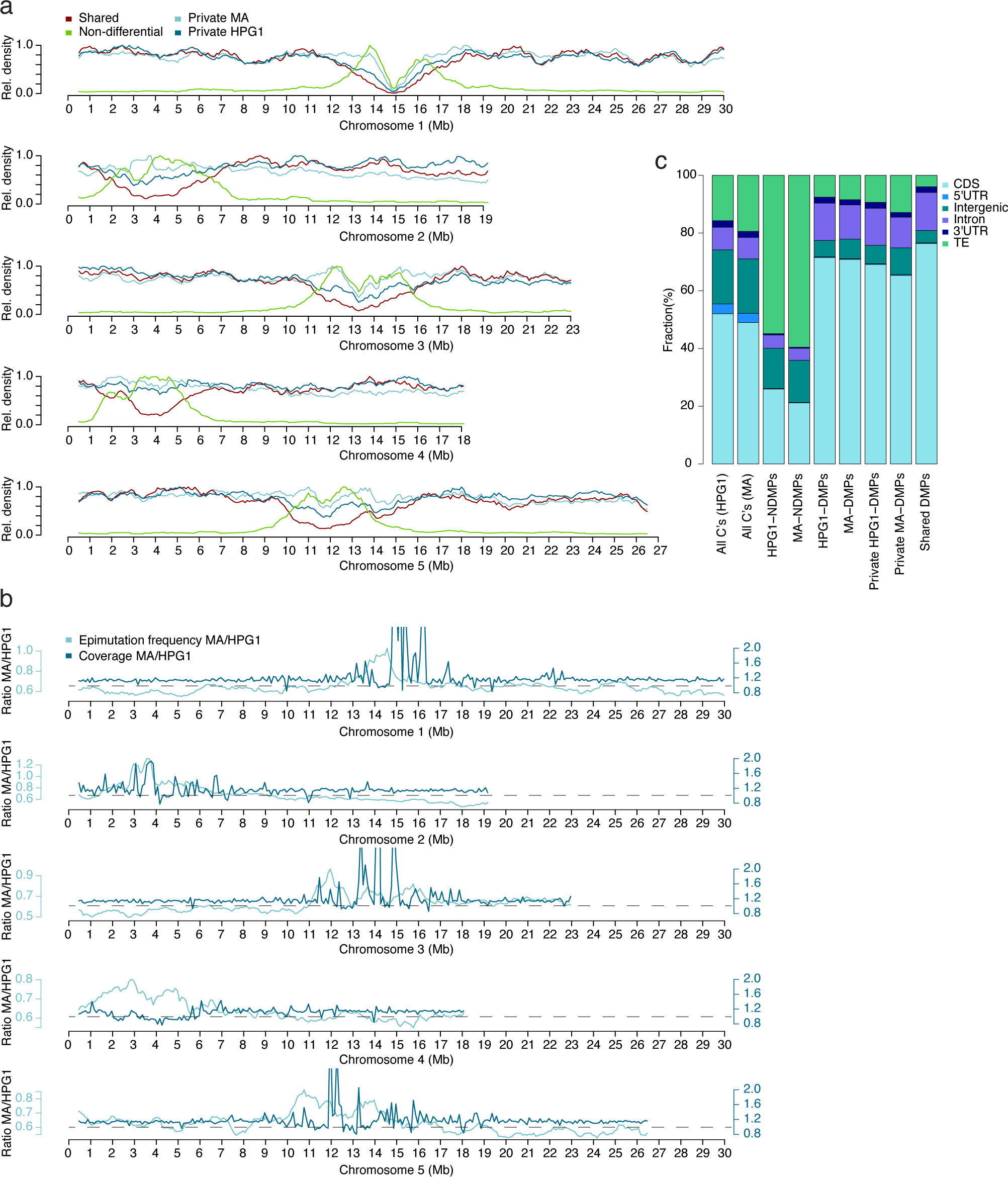
Methylated cytosines independently identified as DMPs in HPG1 accessions and/or MA lines. (a) Relative density of DMPs along the 5 chromosomes. For each class and each chromosome, the window with the maximal density was set to 1. Sliding window; window size 100,000 bp; step size 10,000 bp. (b) Ratios between epimutation frequencies and sequencing depth along the 5 chromosomes for MA and HPG1 lines. Epimutation frequencies were determined as the number of DMPs per cytosine with at least threefold coverage per window. Coverage is represented as average coverage per window across all accessions of each population. Dashed lines mark the ideal coverage ratio of 1. Sliding window; window size 100,000; step size 10,000 bp. **(c)** Annotation of Cs, N-DMPs and DMPs. Cytosines were hierarchically assigned to CDS > intron > 5’ UTR > 3’ UTR > transposon > intergenic.

**Figure S14.**
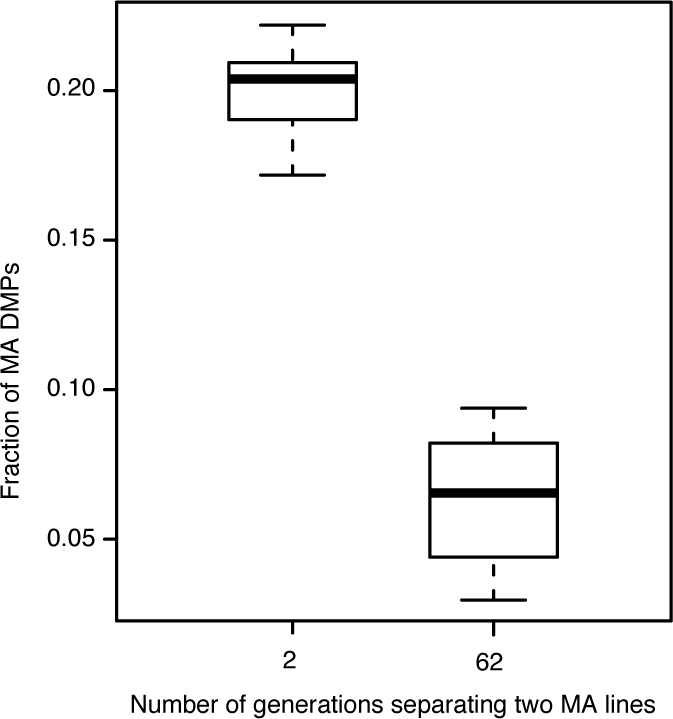
MA DMPs shared with HPG1 DMPs according to MA generational distance. We computed DMPs between two randomly chosen MA strains separated by specific numbers of generations and plotted the fraction of those DMPs shared with a randomly chosen HPG1 strain. Each boxplot summarizes ten such random comparisons.

**Figure S15.**
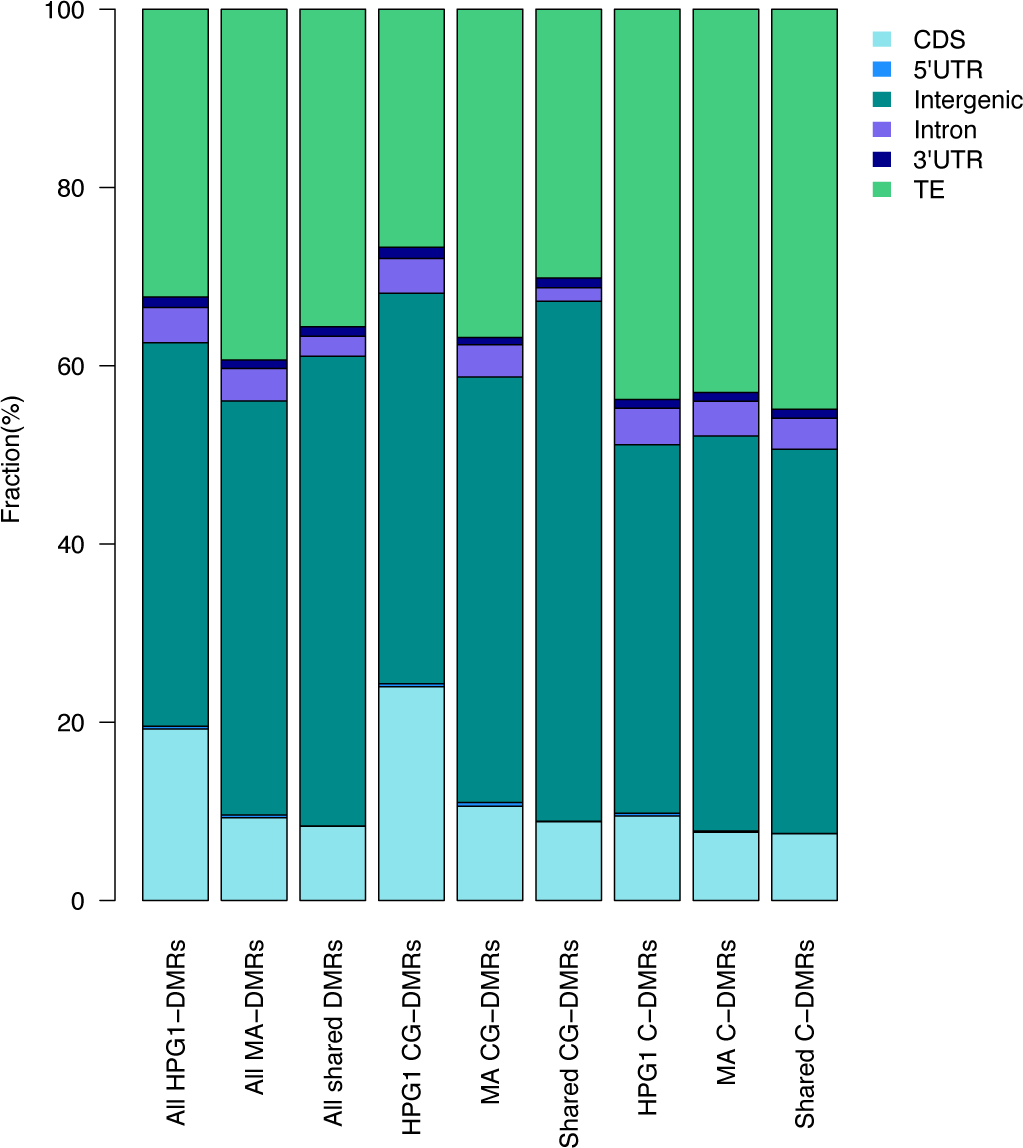
Overlap of MA DMRs and HPG1 DMRs per genomic feature and DMR sequence context. DMRs were hierarchically assigned to CDS > intron > 5’ UTR > 3’ UTR > transposon > intergenic. CG-DMRs had significantly different methylation in the CG context only and C-DMRs in any other (additional) context(s).

**Figure S16.**
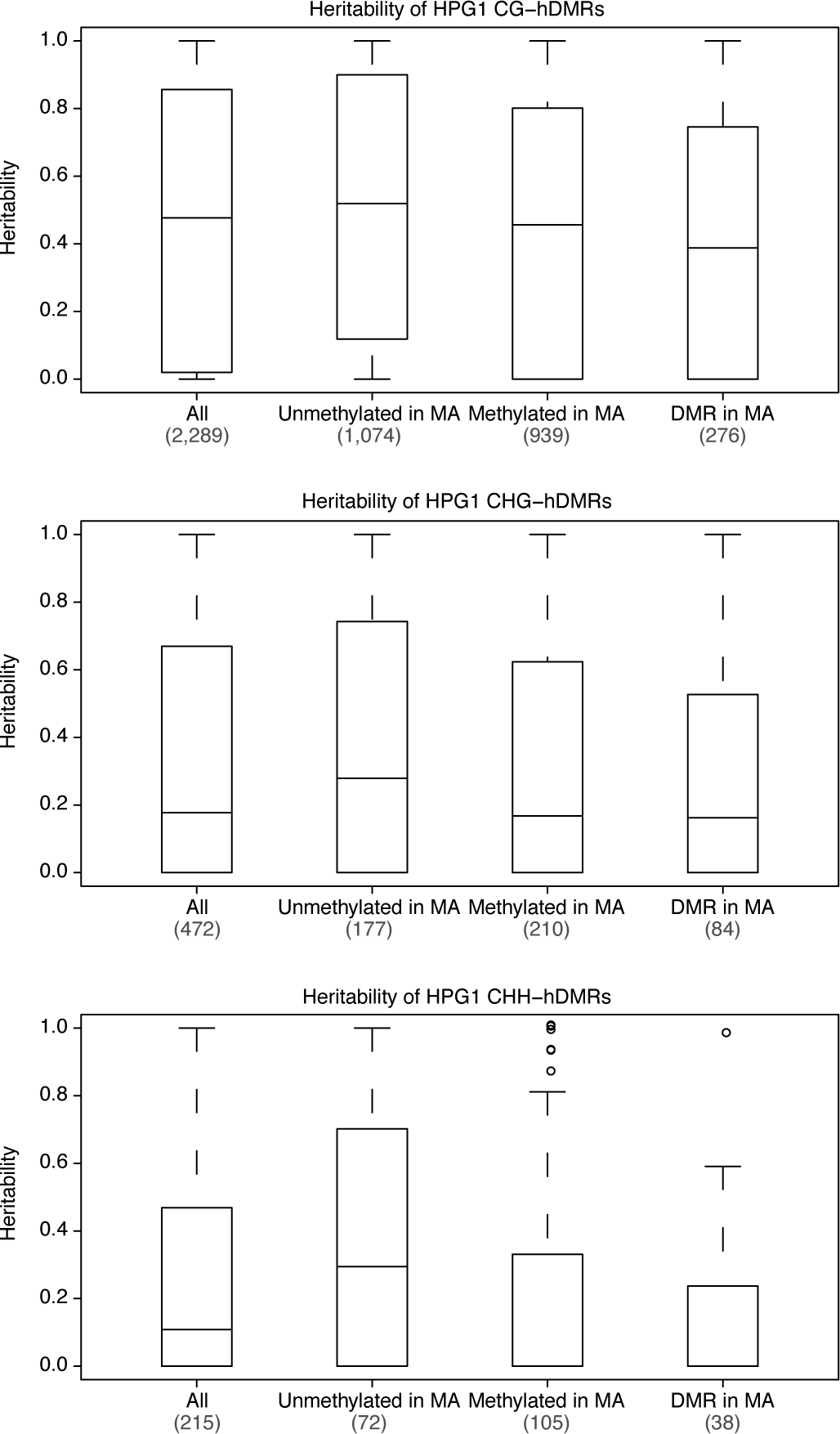
Heritability by sequence context of hDMRs in HPG1, and their overlap with unmethylated, methylated and differentially methylated regions in MA lines.

**Figure S17.**
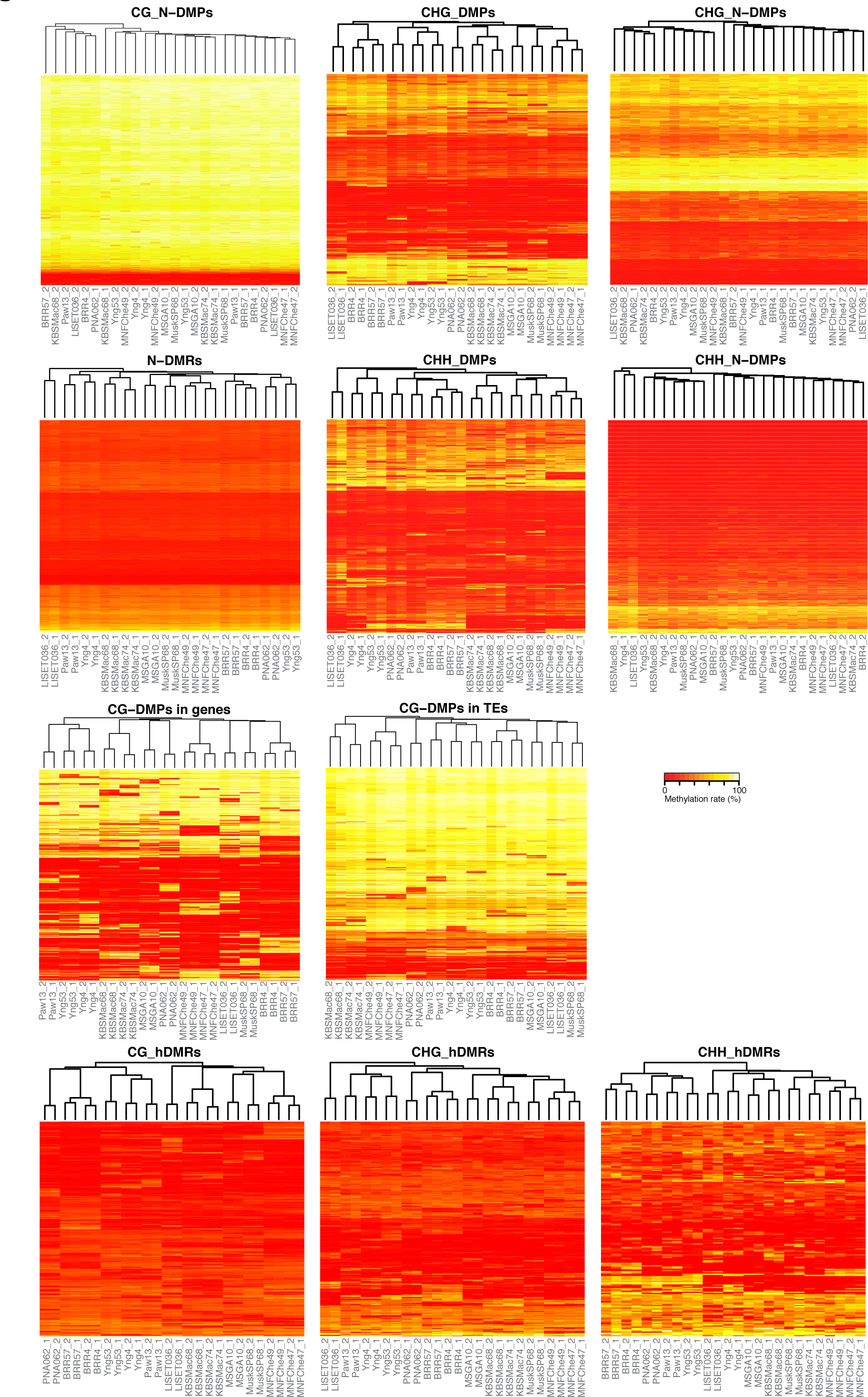
Hierarchical clustering by polymorphic and non-polymorphic positions and regions.

**Figure S18.**
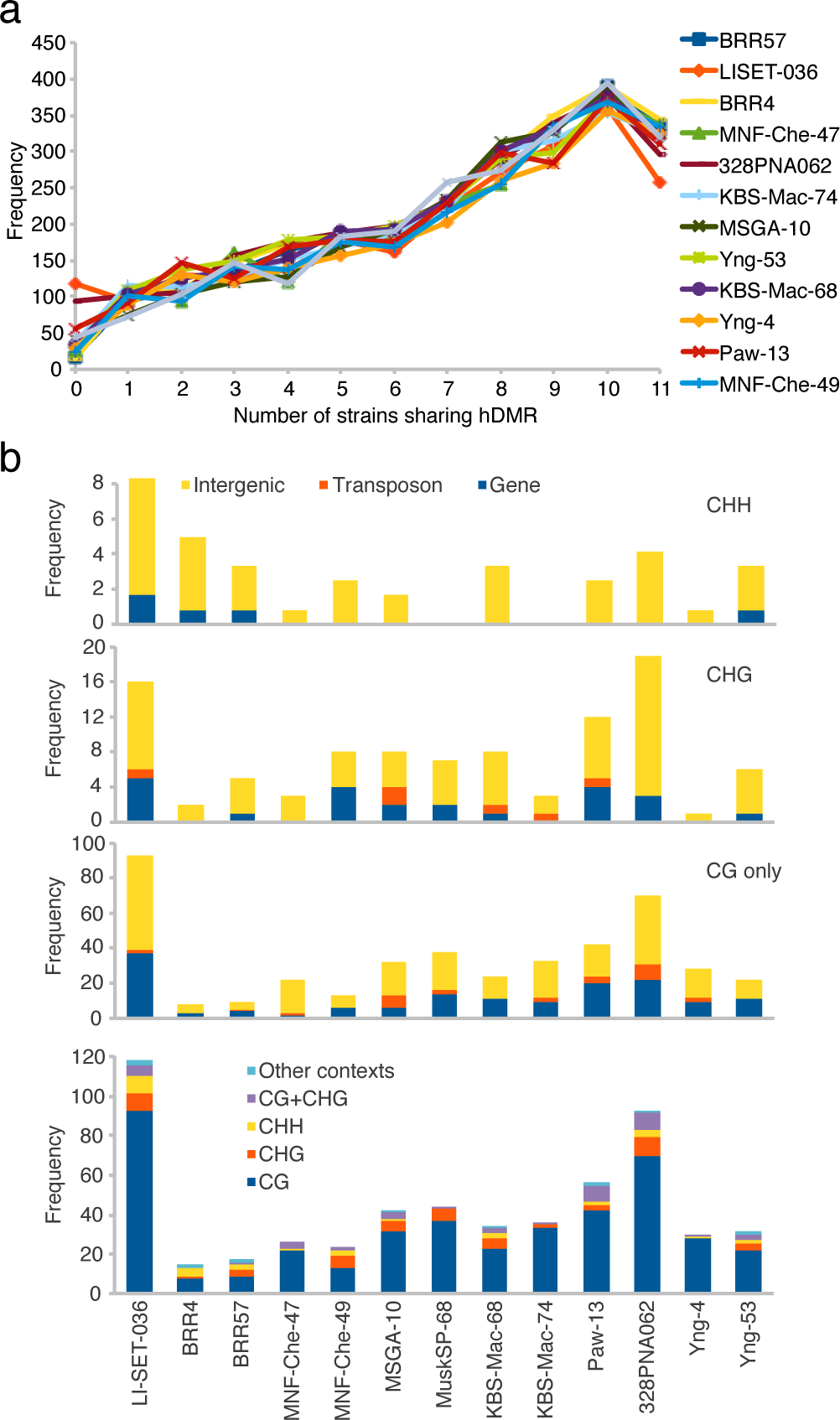
Analysis of LI-SET-036 private hDMRs. (a) The number of strains sharing the same methylation status for DMRs found in each strain is plotted (determined by the strain grouping procedure; see Methods). (b) Stacked bar plots showing the distributions of sequence contexts (bottom) and overlapping genomic features (top three plots) for each strain’s private hDMRs. ‘CG only’ exclusively considers CG-hDMRs whereas ‘CHG’ and ‘CHH’ might also include hDMRs of other contexts than CHG and CHH, respectively. The distribution across intergenic space, TEs and genes was similar for all strains. See section “Analysis of LI-SET-036 specific hDMRs” above for more details.

### Supplementary tables

**Table S1: Haplogroup-1 (HPG1) accessions used in this study.**

**Table S2: Common SNPs and SVs.**

**Table S3: Segregating SNPs and SVs.**

**Table S4: Summary statistics on methylome sequencing.**

**Table S5: Methylated regions (MRs).**

**Table S6: Differentially methylated regions (DMRs).**

**Table S7: Highly differentially methylated regions (hDMRs).**

**Table S8: Summary statistics on transcriptome sequencing.**

**Table S9: Differentially expressed (DE) genes identified in pairwise comparisons between HPG1 accessions.** All DE genes (q-value < 0.05) between any two accessions are listed. If more than one gene name appears in column 1, the read counts could not be assigned to one gene in particular and/or a fused transcript was suggested by the read data.

**Table S10: Statistics of overlapping DE genes and hDMRs.**

**Table S11: Overlap of SVs with MRs in HPG1 and Col-0 in different genomic features.**

**Table S12: Scoring matrices for SNP calling and assessing cytosine site statistics (for bisulfite sequencing).**

